# Sand dropseed (*Sporobolus cryptandrus*) – A new pest in Eurasian sand areas?

**DOI:** 10.1101/2021.07.05.451115

**Authors:** P. Török, D. Schmidt, Z. Bátori, E. Aradi, A. Kelemen, A. A. Hábenczyus, C. P. Diaz, C. Tölgyesi, R.W. Pál, N. Balogh, E. Tóth, G. Matus, J. Táborská, G. Sramkó, L. Laczkó, S. Jordán, J. Sonkoly

## Abstract

For the effective control of an invasive species, gathering as much information as possible on its ecology, establishment and persistence in the subjected communities is of utmost importance. We aimed to review the current distribution and characteristics of *Sporobolus cryptandrus* (sand dropseed), an invasive C4 grass species of North American origin recently discovered in Hungary. We aimed to provide information on (i) its current distribution paying special attention to its invasion in Eurasia; (ii) the characteristics of the invaded habitats in Central Europe; (iii) seed bank formation and germination characteristics, crucial factors in early establishment; and (iv) the effects of its increasing cover on vegetation composition. Finally, we aimed to (v) point out further research directions that could enable us to understand the invasion success of this potential invasive species. Field surveys uncovered large stands of the species in Central and Eastern Hungary with most of the locations in the former, especially the Kiskunság region. The species invaded disturbed stands of dry and open sand grasslands, closed dune slack grasslands and it also penetrates into natural open sand grasslands from neighbouring disturbed habitats. Increasing cover of *Sporobolus cryptandrus* caused a decline in species richness and abundance of subordinate species both in the vegetation and seed banks, but a low density of *Sporobolus cryptandrus* can even have a weak positive effect on these characteristics. Viable seeds of *Sporobolus* were detected from all soil layers (2.5 cm layers measured from the surface to 10 cm in depth), which indicates that the species is able to form a persistent seed bank (1,114 to 3,077 seeds/m^2^ with increasing scores towards higher abundance of the species in vegetation). Germination of *Sporobolus cryptandrus* was negatively affected by both litter cover and 1 cm deep soil burial. To sum up, *Sporobolus cryptandrus* can be considered as a transformer invasive species, whose spread forms a high risk for dry sand and steppe grasslands in Eurasia. We can conclude that for the effective suppression of the species it is necessary: (i) to clarify the origin of the detected populations; (ii) to assess its competitive ability including its potential allelopathic effects; (iii) to assess its seed bank formation potential in habitats with different abiotic conditions; and (iv) to assess the possibility of its suppression by natural enemies and management techniques such as mowing or livestock grazing.

## Introduction

The distribution range and abundance of invasive plants have dramatically increased in recent decades providing a serious challenge for protection, conservation, and restoration of natural and semi-natural habitats worldwide (van Kleunen et al. 2019, Pyšek et al. 2020). While casual establishment of alien species in various natural ecosystems became a relatively frequent phenomenon as a consequence of increased human influence, transformer invasive species form one of the most serious threats for natural communities and ecosystems (Richardson et al. 2000). Transformer invasive species often reduce biodiversity, alter disturbance regimes, and affect ecosystem structure and functions in the subjected communities (Richardson et al. 2000, Byers et al. 2010, Catford et al. 2011).

For the effective control of an invasive species, it is crucial to collect as much information as possible on (i) their ecology, especially establishment and persistence characteristics, and on (ii) communities potentially threatened by its invasion. It is also crucial to detect the plant invasion in an early stage, when the distribution of the invasive species is still limited to one or a few isolated locations, where its eradication still might be possible. However, this is rather challenging for inconspicuous species (such as certain grasses), which are usually difficult to determine and to detect (Jarić et al. 2019). Members of the Poaceae and Asteraceae families contribute most of the aggressive invasive plant species across the globe (Pyšek et al. 2017). Invasions of many short-lived and perennial grasses present serious problems worldwide including grasses characterised either by C3 or C4 photosynthetic pathway (D’Antonio & Vitousek 1992, Fusco et al. 2019, van Kleunen et al. 2019). The C4 photosynthetic pathway provides many advantages over the C3 one in arid and warm climate, such as carbon fixation with a lower water cost, higher temperature optimum for carbon fixation, lower sensitivity to water stress and high fire resistance (Johnston 1996).

Thermophilic neophytes include numerous species of Poaceae with the C4 photosynthetic pathway, which have become constant elements of some warm ruderal communities in Eurasia (Leuschner & Ellenberg, 2017). Introduced C4 grasses have already caused dramatic losses of biodiversity in the Americas (e.g., savanna and forest ecosystems in central and South America or desert grasslands and dry woodlands in the North America, Williams & Brauch 2000) or in Australia (e.g., tropical grasslands in Australia, Brooks et al. 2010). Similarly, the Eurasian steppe zone is characterised by the prevailing dominance of C3 grasses with only some notable exceptions of C4 species (e.g., *Botriochloa ischaemum*, *Cynodon dactylon*), which are thought to be introduced in historical times (Hurka et al. 2019). According to the projected climate change scenarios, global temperatures will increase in the future, likely resulting in an increased expansion of C4 grasses in plant communities of arid environments of Eurasia. Direct and indirect effects of climate change include the increase of minimum and maximum temperatures, the increasing frequency and magnitude of droughts in the vegetation period, and the changing annual distribution of precipitation shifting the peak of precipitation from the vegetation period to the dormant period (in Central Europe shifting from the summer period to winter, IPCC 2013). This means that increased levels of aridity, increased likeliness of weather extremities, and the associated increased risk of extreme fire events favour the formation and spread of more drought-adapted communities and species. The decline of dominant native C3 grasses like *Festuca* species increases the risk of invasion via the colonization of drought-adapted non-native C4 species. The effects of climate change may be amplified at the regional scale by large-scale water regulation works and the increased demand for irrigation in agricultural areas, or by the high transpiration rate of established non-native tree plantations (Tölgyesi et al. 2020). The water-stressed open sand grasslands are excellent targets for the establishment of non-native C4 plant species. The establishment and the effects of several invasive herbaceous species on the native communities have been studied and reported for sand regions (e.g., *Asclepias syriaca* – Kelemen et al. 2016 or *Conyza canadensis* – Mojzes et al. 2020).

In the current paper, we aim to study the current distribution and characteristics of *Sporobolus cryptandrus* (sand dropseed), an invasive C4 grass species of North American origin, recently detected in Hungary (Török & Aradi 2017). We aim to evaluate its effects on the native sand grassland vegetation by analysing stands along an increasing *Sporobolus* abundance gradient. In particular, we aim to provide information on (i) the current distribution of the species with special attention to its invasion in Eurasia by summarising published occurrence data, (ii) the characteristics of the invaded habitats in Central Europe, (iii) seed bank formation and germination characteristics, crucial in early establishment of the species, and on (iv) the effects of increasing cover of the species on vegetation composition. Finally, we aim to (v) point out further research directions that would help to understand the invasion success of this species, and to (vi) evaluate possible management techniques to control it.

## Materials and methods

### Morphological characteristics and ecology of the species

The species is a member of the dropseed genus (*Sporobolus* R.Br.) consisting of more than 160 species with the highest number of endemic species in Africa, Australasia, North and South America (Simon & Jacobs 1999, Király & Hohla 2015). Species in the genus are typical in tropical and warm temperate climate, generally tolerate drought but can also be found in saline habitats from loose sandy soils to heavy floodplain soils (Simon & Jacobs 1999). Sand dropseed (*Sporobolus cryptandrus* (Torr.) A.Gray) is a perennial bunchgrass with a height of 40–80 cm (up to 100 cm with inflorescences). Both the auricula and the ligula are very short and at the orifice of the sheaths, on the leaves’ margin around the nodes there is a collar of dense white hairs, but scattered hairiness is typical also for the whole leaf edge. The edge of the 4–5 mm wide leaves is sharp, but the leaves are softer than the leaves of a *Calamagrostis*. The inflorescences are at least partly covered by the flag leaf but are very similar to those of an *Agrostis*, not surprisingly formerly the genus was classified together with the latter one (Figure 1, Simon & Jacobs 1999). The species is characterised by a C4 photosynthetic pathway. It produces very tiny propagules in high abundance (the caryopsis is *ca* 1 mm in length); based on literature data, one individual is able to produce up to ten-thousand seeds (Brown 1943). The thousand-seed weight of the species is 0.083 g (Török et al., unpublished). It was also reported that the epicarp of the seeds becomes sticky when wet, which, besides the small weight of the seeds, may contribute to its effective dispersal (Holub & Jehlík 1987). Seeds have high viability but a high proportion of them can become dormant, which means that dormancy breaking in the form of stratification, moist conditions and/or scarification is necessary for successful germination (Holub & Jehlík 1987, Sartor & Malone 2010). The species is likely able to build up persistent seed banks in its native range (Clements et al. 2007) and might have an allelopathic effect on the germination of other species as extrapolated from *Sporobolus pyramidatus* (Lam.) Hitchc. (Rasmussen & Rice 1971).

**Figure 1.**
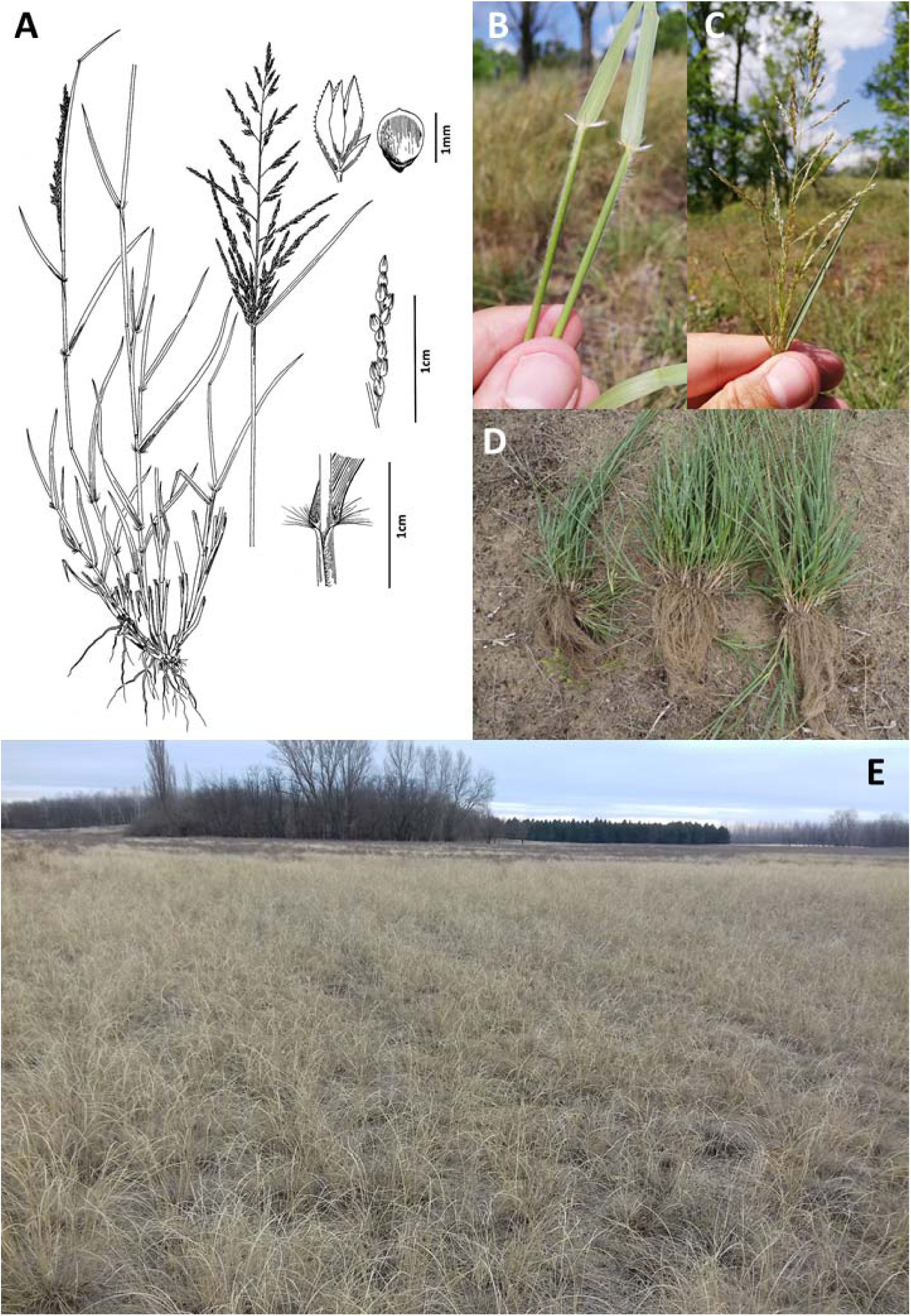
The habitus and morphological characteristics of *Sporobolus cryptandrus*. Notations: A) habitus and morphological characteristics of the species (drawings by J. Táborská), B) Nodes with leaves and C) inflorescences (photos by Z. Bátori); D) root system of the species (photo by E. Aradi); and E) a site in the Kiskunság region with a mass invasion by *S. cryptandrus* (photo by C. Tölgyesi).

### Distribution range of the species

The native range of *S. cryptandrus* lays in North America including the United States, Southern Canada, and the northern part of Mexico (Britton & Brown 1970; Holub & Jehlík 1987; Lackschewitz 1991; Nobis et al. 2015). The species is typical in short-grass prairies, sagebrush deserts, and chaparral communities but sometimes also enters the sagebrush steppe (Hitchcock et al. 1969, Tilley et al. 2009, Lesica 2012). In its native range it is a member of the climax plant communities on deep sands, while on heavier soils it is an early successional colonizer. The plant is extremely drought-tolerant and it is highly competitive with co-occurring native species even in desert climates (Wan et al. 1993, Ogle et al. 2009, Tilley et al. 2009). Typically, it grows at lower elevations on sandy soils (which explains the English colloquial name: sand dropseed), mainly on disturbed sites such as dry riverbeds, rocky slopes, and along roadsides. It can also be found at higher elevations and coarser soils. The plant is extremely drought tolerant and highly competitive with co-occurring native species even in desert climates (Wan et al. 1993, Ogle et al. 2009, Tilley et al. 2009).

Outside of its native range, the species has been reported from Australia and Tasmania, Japan, New Zealand, and Argentina (Edgar & Connor 2000, Curto 2012, Randall 2017). In Eurasia, the species was detected formerly in several locations (Figure 2). *Sporobolus cryptandrus* is known from isolated locations from Austria, France, Germany, Italy, the Netherlands, Russia, Slovakia, Spain, Switzerland, Ukraine, and the United Kingdom (Murr 1902, Thellung 1919, Ryves et al. 1988, Sani et al. 2015, Dflor 2021, NBMC 2021, Electronic Appendix 1). Amongst the first naturalised populations was a riverbank near Bratislava, Slovakia (Holub & Jehlík 1987). Large-scale spreading was reported also into steppe habitats in Western Russia and Ukraine (Alekseev et al. 1996, Kuvaev & Stepanova 2014, Demina et al. 2016, Gouz & Timoshenkova 2017, Demina et al. 2018, Maltsev & Sagalev 2018). Historical data of the species were reported from the western part of Hungary, near to the city of Győr (= *Sporobolus subinclusus*, 1927 in Polgár 1933, one specimen detected from the territory of an oil seed factory, and likely originated from Argentina), but the data cannot be validated and was not supported with a herbarium sheet. In 2016, the species was discovered in two sandy regions of Hungary, in the city of Debrecen (Nyírség region, acidic sand) and near the town Kiskunhalas (Kiskunság region, calcareous sand) in several small locations (Török & Aradi 2017, Erdős et al. 2018, Molnár et al. 2020).

**Figure 2.**
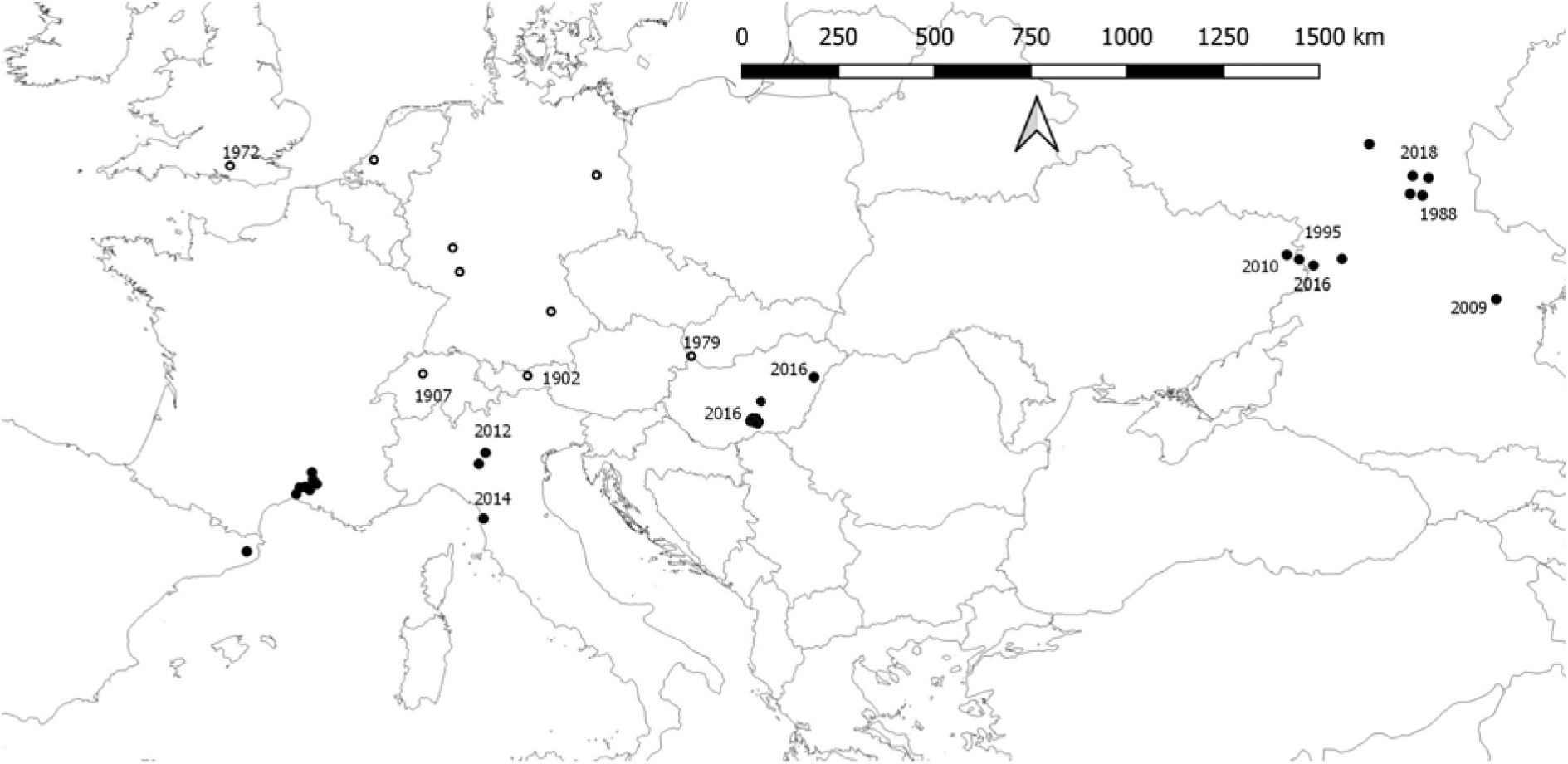
The distribution of *Sporobolus cryptandrus* in Eurasia. The full symbols show naturalised populations, while the empty ones denote casual establishment. See more details on locations in Electronic Appendix 1.

### Vegetation and soil sampling

After its discovery in Eastern and Central Hungary (Debrecen, Nyírség region and Kiskunhalas, Kiskunság region, respectively; Török & Aradi 2017) more detailed and systematic surveys were initiated. The largest locality of the species in Debrecen and four localities with large established populations in the Kiskunság region were selected for a detailed vegetation sampling (Table 1). In each site, plots along an increasing cover gradient of *Sporobolus cryptandrus* were sampled. We sampled reference plots with no *Sporobolus* (cover category I), and plots characterised with 1-25%, 26-50% or 51-75% cover of the species (cover categories II, III and IV, respectively), but in the latter case *Sporobolus* cover rarely exceeded 70%. Altogether, 10 plots per cover group (altogether 40 plots per site) were recorded, the percentage cover of all vascular plant species were assessed in the summer of 2019.

**Table 1.** Sample sites in Central and East Hungary.

| Sample sites | Sample code | Nearest town | GPS coordinates |  | Habitat type | Population size of *Sporobolus* |
| --- | --- | --- | --- | --- | --- | --- |
| Debrecen | D | Debrecen | 47.55422 N | 21.61537 E | Degraded sand grassland in urban area | 4,000-6,000 |
| Kiskunhalas North | KN | Kiskunhalas | 46.48243 N | 19.47987 E | Mosaic of natural sand steppe and pine plantations | 5,000-7,000 |
| Katonatelep | KT | Kecskemét | 46.95044 N | 19.76217 E | Grazed meadow-steppe | 200,000-300,000 |
| Airport | A | Kecskemét | 46.92774 N | 19.72590 E | Sandy old-field (30+-year-old) | 75,000-100,000 |
| Kiskunhalas East | KE | Kiskunhalas | 46.40935 N | 19.56305 E | Disturbed sandy forest-steppe | 1,500-2,000 |

For detailed soil analyses, we sampled the topsoil (<5 cm) with a small spade from 40 random locations per site near the vegetation plots (10 samples for each cover category pooled, about 500 g air-dried soil per pooled sample, 4 pooled samples per site). The following soil characteristics were measured: pH (KCl), soil compactness (this figure is strongly related to the physical texture of the soil; higher scores refer to higher proportion of loam-clay), calcium – CaCO_3_ (m/m%), humus (m/m%), nitrogen – NO_2_ + NO_3_ content (mg/kg), phosphorous – P_2_O_5_ (mg/kg), potassium – K_2_O (mg/kg). Soil analyses were conducted in an accredited laboratory (SYNLAB, Mosonmagyaróvár, Hungary) based on the standardised methods included in the Hungarian standards MSZ-08-0205:1978 (Evaluation of some chemical properties of the soil. Laboratory tests) and MSZ-08-0206-2:1978 (Determination of physical and hydrophysical properties of soils).

### Soil seed banks

We screened the composition of soil seed banks in plots characterised by different levels of *Sporobolus* cover at the Debrecen site. In three plots per cover group, we collected 10 soil cores (10 cm depth and 2 cm diameter) separated to four vertical segments (0–2.5 cm, 2.5–5 cm, 5–7.5 cm and 7.5–10 cm) in the last week of August 2020. Identical vertical segments were pooled per plot. We used the seedling emergence method with bulk reduction by ter Heerdt et al. (1996). Concentrated samples were spread on the surface of pots filled with steam-sterilised potting soil. The samples were regularly watered and checked for emerged seedlings. Seedlings were identified and removed; unidentified seedlings were transplanted and grown until the final identification. Emergence lasted about 11 weeks in the autumn of 2020, from 28^th^ August until 15^th^ November. At the end of the germination period, we identified all seedlings at the highest possible taxonomic level. As we conducted a preliminary germination experiment in the spring, we were able to distinguish the seedlings of *Sporobolus cryptandrus* from the seedlings of native C3 grasses at a very early stage. Other C4 grasses typical in the region were almost absent from the plots (e.g., *Cynodon dactylon*) or were present with low density only in seed banks and flowered already very early in the pots (*Eragrostis minor*). Only a small fraction of the seedlings perished before identification to a respective family, genus or species (8 individuals, less than 0.4% of all seedlings, omitted from analyses). Altogether 28 taxa were identified at the species level. We were not able to identify non-septate *Juncus* (1 seedling, treated as *J. conglomeratus/effusus*), or *Epilobium* seedlings (3 seedlings, *Epilobium* sp.) on the species level, and we also pooled the seedlings of *Arenaria leptoclados* and *A. serpyllifolia* as *A. leptoclados*. Seedlings of short-lived small *Veronica* species (*V. polita, V. triphyllos, V. verna*) were pooled as *Veronica* sp. (altogether 65 individuals). For some graminoid seedlings we were able to identify them only at the family level – Poaceae (altogether 38 individuals – these were most likely the seedlings of *Poa angustifolia* or *Lolium perenne*).

### Germination experiment

A greenhouse experiment was conducted to test the effects of increasing seed burial depth (0, 0.5 and 1 cm soil) and increasing levels of litter cover (0, 150 and 300 g/m^2^) and their interaction on the germination potential of *S. cryptandrus* in a full-factorial design with nine treatments in five replications (resulting in 45 pots total). For the selection of the soil and litter cover thicknesses we used a modified version of the experimental setup published by Sonkoly et al. (2020). *S. cryptandrus* seeds were collected in 2019 and dry-stored at room temperature (20–25 °C) in the seed collection of the Department of Ecology at the University of Debrecen. A total of 45 pots filled with steam-sterilized potting soil were used in the experiment and 25 *Sporobolus* seeds were spread out evenly on the surface of each pot (in total 1125 seeds were sown). We used the same sterilized potting soil and the litter of *Festuca rupicola* for covering the seeds. The germination experiment lasted ten weeks from 26^th^ March until 27^th^ May 2020. The seedlings were regularly counted and removed. Only those seedlings were counted which appeared at the surface of the treatment. We registered the number of established seedlings weekly and removed and registered the perished ones – so we were able to calculate the seedling survival rates for the entire experiment.

### Statistical analyses

We calculated species richness (S, number of species), Shannon diversity (H, based on ln) and evenness scores (following Pielou (1975) where Evenness = H/log(S), and H refers for Shannon diversity and S for species richness) to compare vegetation and seed bank data of plots with increasing cover of *Sporobolus*. The effect of site, *Sporobolus* cover and their interaction on the selected vegetation characteristics were analysed using two-way ANOVA, where the fixed factors were ‘sampling site’ and ‘*Sporobolus* cover’, and dependent variables were species richness, Shannon diversity, and evenness with and without the inclusion of *Sporobolus* and its abundances. The effect of litter cover, soil burial and their interaction on the number of germinated seedling and survival rates were analysed by two-way ANOVA, where the fixed factors were ‘litter cover’ and ‘soil burial depth’. The effect of ‘*Sporobolus* cover’ and ‘soil layer’ (included as fixed factors) on the species richness, Shannon diversity and evenness (dependent variables) were analysed by a two-way ANOVA.

The vegetation composition of the five sites were compared with a PCA ordination, where the main data matrix was the species composition of the plots with different *Sporobolus* cover (at each mass locality site. Altogether 10 plots of the same *Sporobolus* cover group were pooled, each cover group was represented by one pooled plot, in total four pooled plots per site), and the secondary matrix contained the selected soil parameters (one pooled sample per *Sporobolus* cover group, in total four pooled samples per site, the means of soil characteristics are shown in Table 2).

**Table 2.**
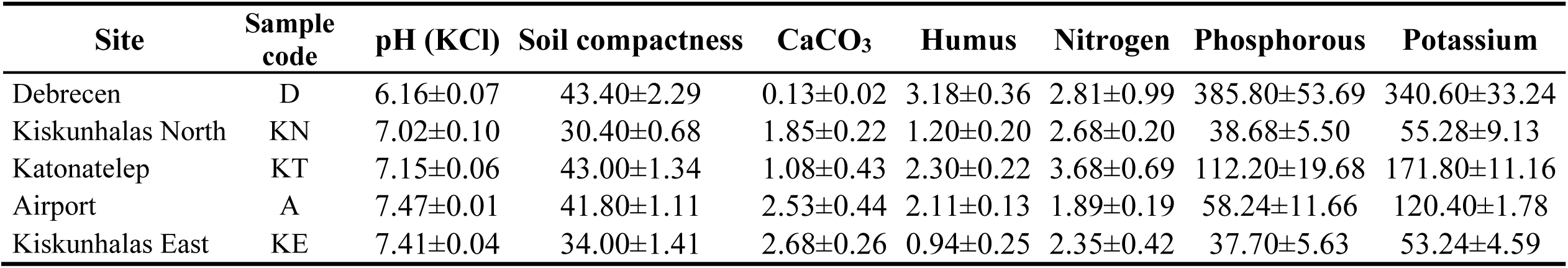
Soil characteristics of the study sites (mean±SE). Measured units: pH (KCl), calcium – CaCO_3_ (m/m%), humus (m/m %), nitrogen – NO_2_ + NO_3_ content (mg/kg), phosphorous – P_2_O_5_ (mg/kg), potassium – K_2_O (mg/kg). Soil compactness is strongly related to the physical texture of the soil; higher scores refer for higher proportion loam-clay fine soil components (e.g., some physical soil texture types are the following: sand = 25-30, sandy loam = 31-37, loam = 38-42, clay-loam = 43-50).

## Results

### Habitat preference of Sporobolus cryptandrus in Hungary

During the detailed field surveys more than 620 individual locations of the species were detected in Central and East Hungary, with most of the locations in the Kiskunság region. In the city of Debrecen, the species was detected in variously degraded and frequently mown urban grasslands situated between blocks of flats, or at road verges, parking lots, and tramlines. In contrast to the urban localities in Debrecen, in the Kiskunság region all *Sporobolus* stands were found in rural landscapes. A remarkable amount of the detected *Sporobolus* populations occur in dry sandy habitats, mainly in disturbed or strongly degraded stands of open sand grasslands, or in closed, desiccated interdune grasslands which typically originate from more wet interdune *Molinia* meadows. Large populations were found along artificial linear landscape elements, such as dirt roadsides, motocross trails, and ploughed fire buffer zones. Furthermore, the species colonised sandy areas ploughed recently (edges of young tree plantations) or during the last three decades. We also found the species in old-fields of various age, both young old-fields characterized by short-lived weeds (e.g., *Anthemis ruthenica, Ambrosia artemisiifolia* or *Bromus tectorum*) and old old-fields already dominated by perennial grasses (e.g., *Festuca pseudovina*, *Cynodon dactylon and Bothriochloa ischaemum*). Further land use types with intense disturbance were also found to facilitate the spread of *Sporobolus cryptandrus*, as we found populations in grasslands formerly or recently overgrazed by sheep, and in the close vicinity of game feeders. Invasion of *Sporobolus* was also detected in grasslands that were burned 10–20 years ago, and successfully regenerated since then (apart from the presence of *Sporobolus*) and consist of the species of natural sandy grasslands (e.g., *Festuca vaginata*, *Koeleria glauca*, *Stipa pennata*, *Alkanna tinctoria*, *Dianthus serotinus and Silene otites*). Moreover, *Sporobolus cryptandrus* spreads in drought-affected stands of open sandy grasslands co-dominated by *Festuca vaginata* and *Stipa pennata*. The southern slopes of sand dunes are typical drought-affected habitats where the destruction of the *Festuca vaginata* tussocks is typical due to severe droughts, and the *Sporobolus* can take its place. *Sporobolus* appeared in the dried-out stands of closed interdune grasslands and in the northern part of the Kiskunság (near Kecskemét) we found it also in the dried-out meadow steppes (with meadow soils). Although primary incursion of *Sporobolus* into natural habitats has not been detected, we found that it can spread to natural open sandy grasslands from the neighbouring disturbed, invaded habitats, where it potentially threatens rare sandy species such as *Dianthus diutinus*, which is a priority species of European community interest. An important observation is that the shade-tolerance of *Sporobolus cryptandrus* is rather low; in the shaded parts of the invaded patches only sparse population of *Sporobolus* were found.

### Soil and vegetation characteristics of the selected sites

Soil analyses of the five study sites revealed that the soil pH ranged between 6.16-7.41, the lowest score in Debrecen (acidic sand deposits) and higher scores were typical in the sample sites of the Kiskunság region (calcareous sand deposits). The physical soil texture type expressed by soil compactness ranged from sand to clay loam, with some sites with a higher load of nutrients, especially in phosphorous (D and KT sites) and potassium (D, KT, and A sites) (Table 2).

Studying the selected five sites, we found that both increasing *Sporobolus* cover, and the sampling site significantly affected most of the studied variables of vegetation with or without the inclusion of the *Sporobolus* cover in the calculations (Table 3). In all sites, we detected an increase in species richness, Shannon diversity and evenness scores from cover group I to II and then a rapid decline was detected for all variables (Figures 3, 4, and 5). There were, however, highly site-dependent differences in the magnitude of this effect, but in most cases no interaction between *Sporobolus* cover and sampling site was detected (Table 3). This pattern was especially distinct for Shannon diversity and evenness scores (Figures 4 and 5). These trends were clearly shown also on the ordination diagram (Figure 4). The ordination clearly revealed that sites with markedly different vegetation composition will be “homogenised” by the increase of the cover of *Sporobolus*, and almost all other species were negatively affected by the high cover of the invasive species (Figure 6). Detailed vegetation compositional data of plots on which Figure 6 was based is summarised in Electronic Appendix 2.

**Figure 3.**
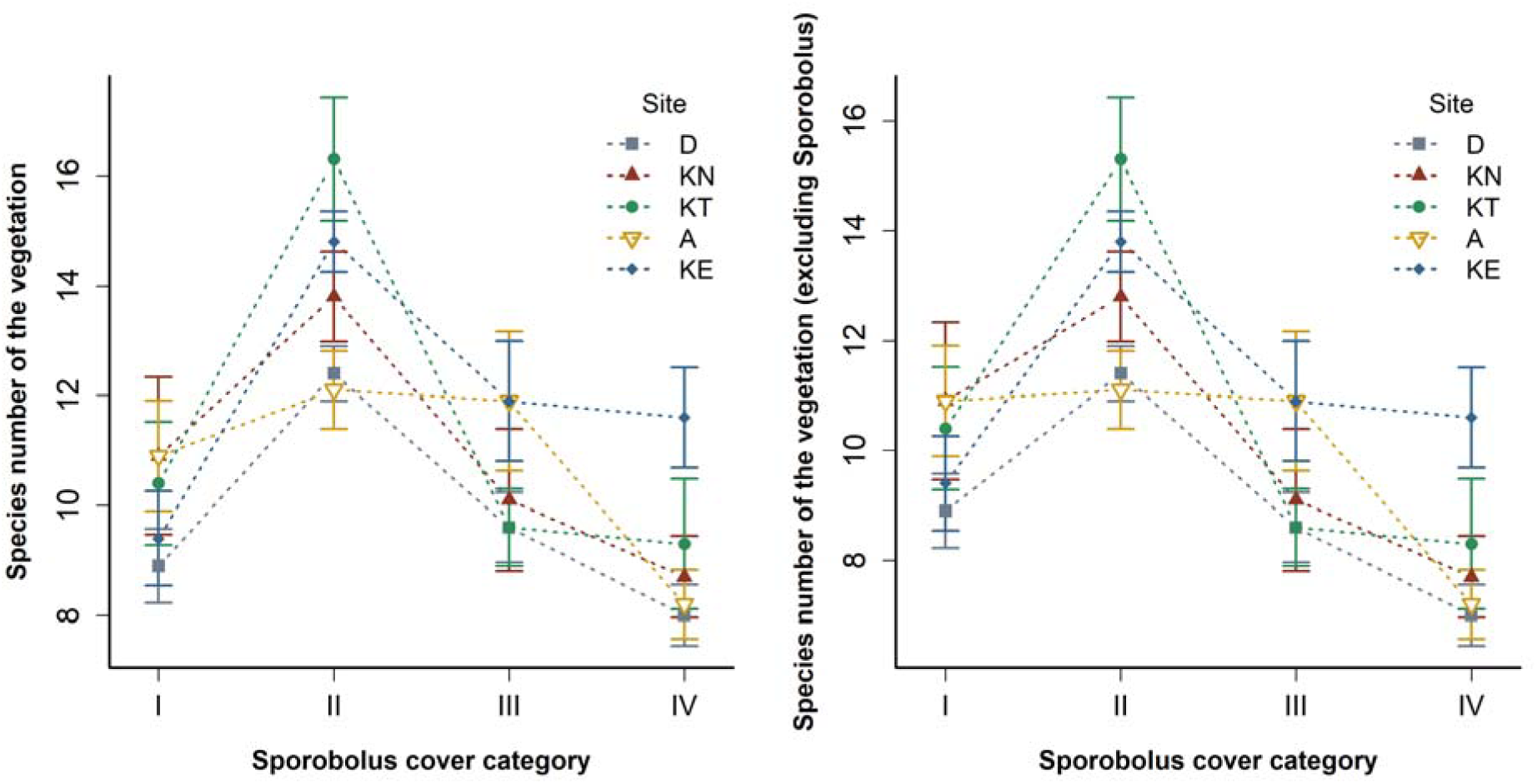
The relationship between *Sporobolus* cover categories and the species richness of the vegetation in the study sites.

**Figure 4.**
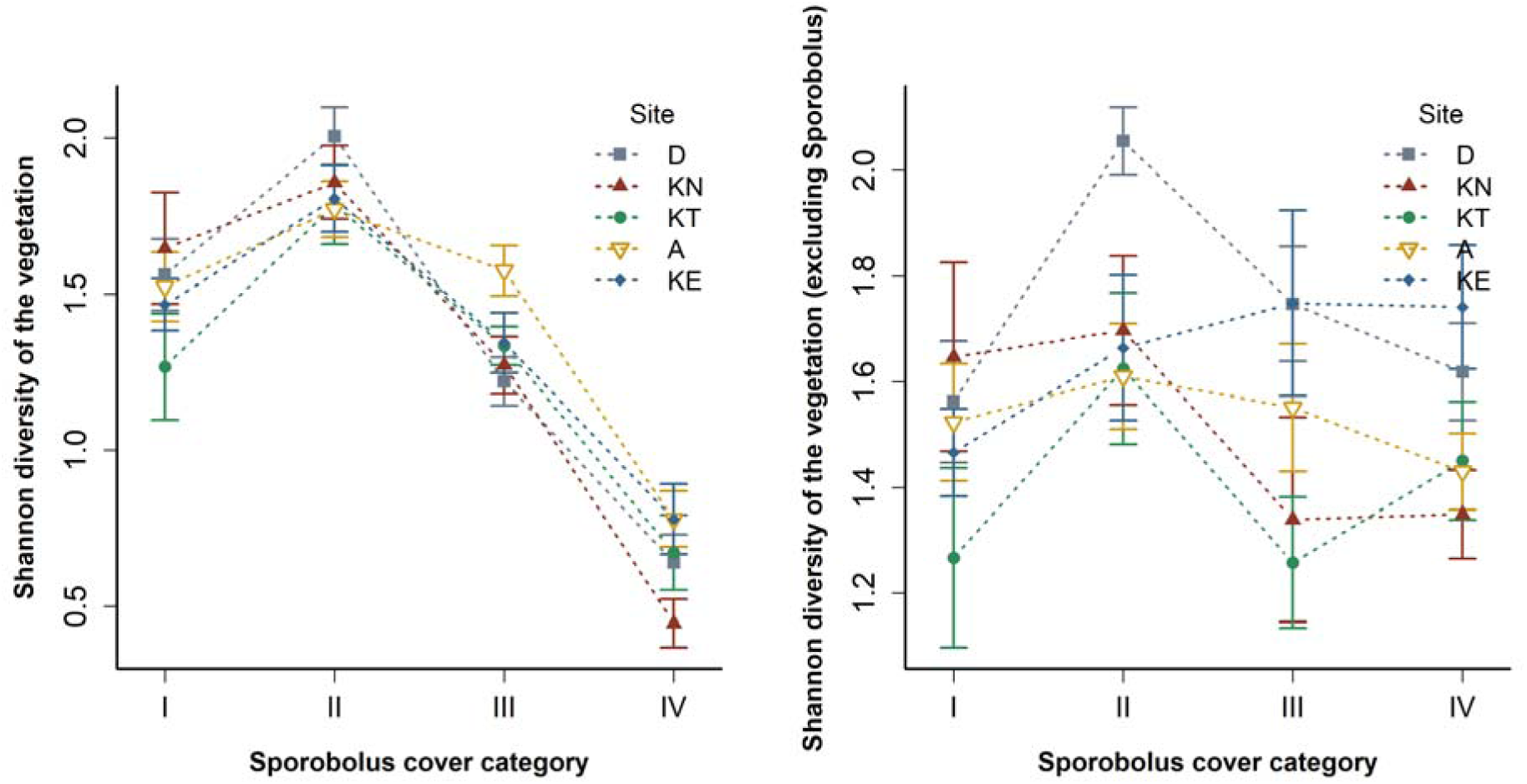
The relationship between *Sporobolus* cover categories and the Shannon diversity of the vegetation in the study sites.

**Figure 5.**
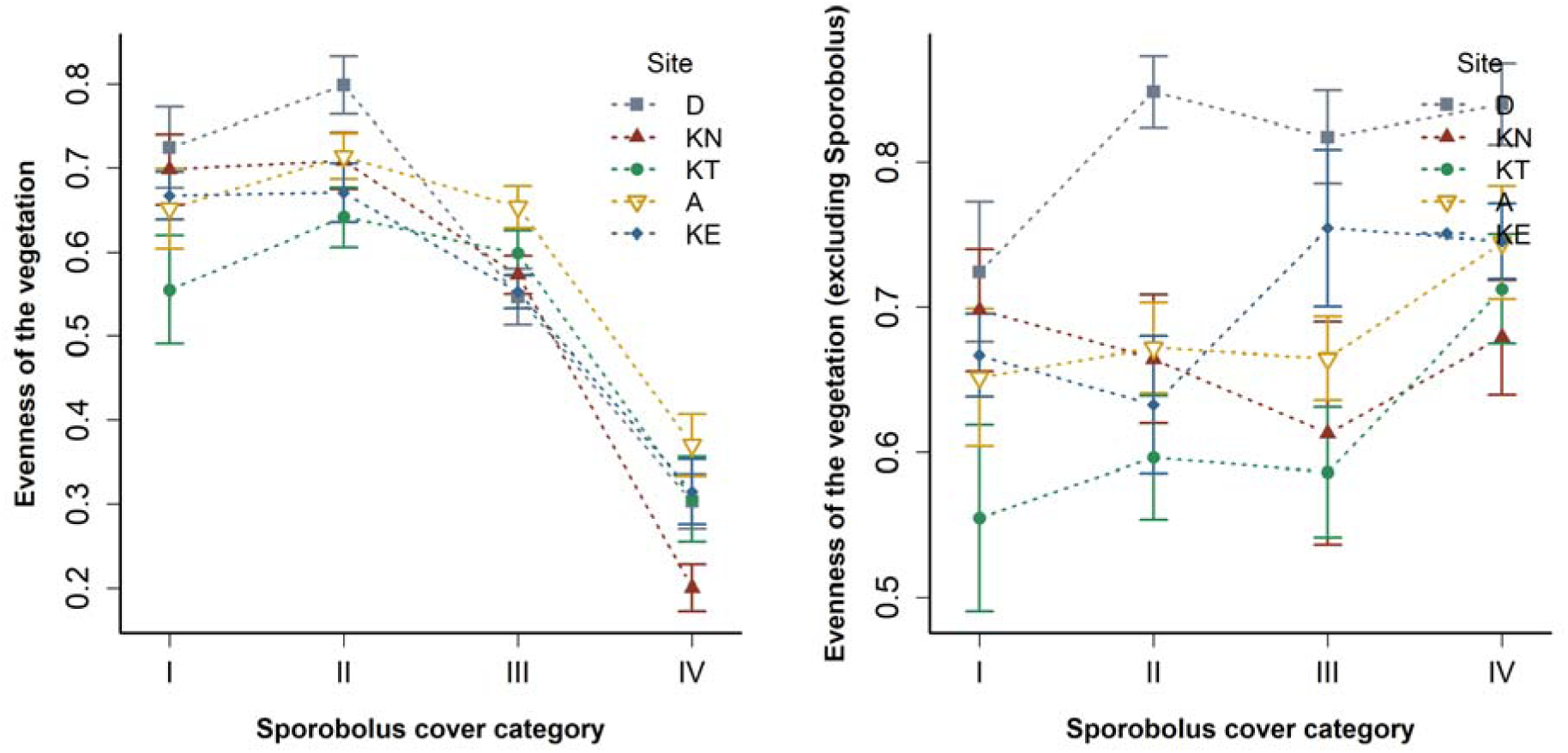
The relationship between *Sporobolus* cover categories and the evenness of the vegetation in the study sites.

**Figure 6.**
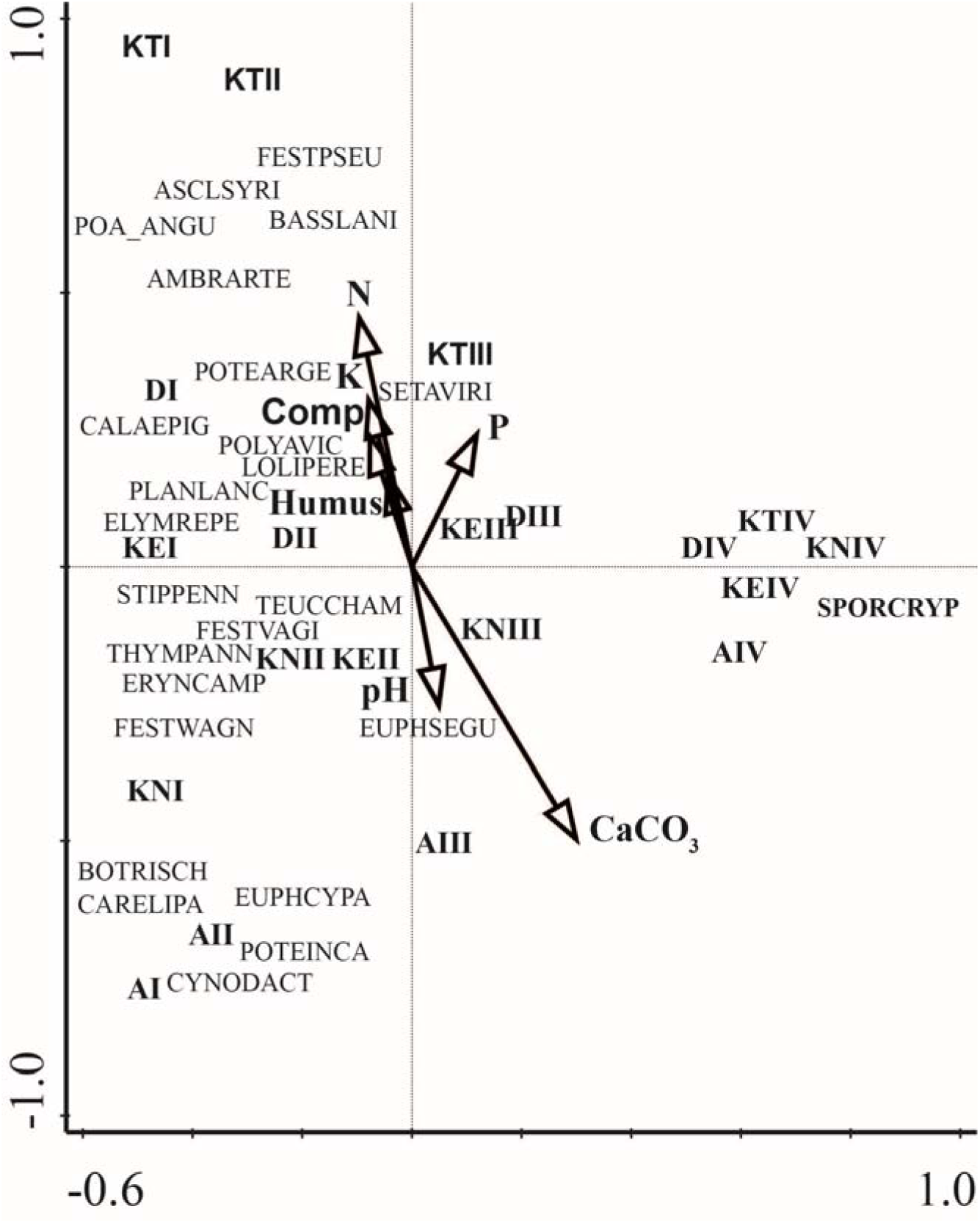
PCA triplot with the 25 most abundant species (Eigenvalues are 0.772 and 0.070 for the first and second axis, respectively). Vegetation composition of the mass locality sites with different cover of *Sporobolus*. Main matrix is the species abundances (10 plots per site and Sporobolus cover categories are pooled, four pooled plots per site are included (I-IV). Sites: D = Debrecen site, KN = Kiskunhalas North, KT = Katonatelep, A = Airport, KE = Kiskunhalas East. Species are abbreviated using the first four letters of the genus names and four letters of the species names. Species are the following: SPORCRYPT = *Sporobolus cryptandrus*, CYNODACT = *Cynodon dactylon*, BOTRISCH = *Bothriochloa ischaemum*, CARELIPA = *Carex liparocarpos*, FESTPSEU = *Festuca pseudovina*, POA_ANGU = *Poa angustifolia*, POTEINCA = *Potentilla incana*, FESTWAGN = *Festuca wagneri*, BASSLANI = *Bassia laniflora*, EUPHCYPA = *Euphorbia cyparissias*, FESTVAGI = *Festuca vaginata*, EUPHSEGU = *Euphorbia seguieriana*, STIPPENN = *Stipa pennata*, POLYAVIC = *Polygonum aviculare*, PLANLANC = *Plantago lanceolata*, ASCLSYRI = *Asclepias syriaca*, ERYNCAMP = *Eryngium campestre*, LOLIPERE = *Lolium perenne*, POTEARGE = *Potentilla argentea*, ELYMREPE = *Elymus repens*, THYMPANN = *Thymus pannonicus*, TEUCCHAM = *Teucrium chamaedrys*, AMBRARTE = *Ambrosia artemisiifolia*, CALAEPIG = *Calamagrostis epigeios*, SETAVIRI = *Setaria viridis*.

**Table 3.** Effect of increasing cover of *Sporobolus cryptandrus* on vegetation characteristics of the subjected plots (two-way ANOVA, significant values are indicated with boldface, p<0.05).

| Vegetation | *Sporobolus* cover |  | Sampling site |  | *Sporobolus* cover ×<br>Sampling site |  |
| --- | --- | --- | --- | --- | --- | --- |
|  | *F*<sub>3,199</sub> | *p* | *F*<sub>4,199</sub> | *p* | *F*<sub>12,199</sub> | *p* |
| **including *Sporobolus*** |  |  |  |  |  |  |
| Species richness | **24.241** | **<0.001** | **3.100** | **0.017** | 1.795 | 0.052 |
| Shannon diversity | **105.361** | **<0.001** | 1.068 | 0.374 | 1.547 | 0.111 |
| Evenness | **121.418** | **<0.001** | **2.898** | **0.023** | **2.280** | **0.010** |
| **excluding *Sporobolus*** |  |  |  |  |  |  |
| Species richness | **22.455** | **<0.001** | **3.100** | **0.017** | 1.795 | 0.052 |
| Shannon diversity | **3.711** | **0.013** | **4.492** | **0.002** | 1.209 | 0.280 |
| Evenness | **3.480** | **0.017** | **10.972** | **<0.001** | 1.079 | 0.380 |

### Seed banks

Altogether 2,132 seedlings of 32 taxa were germinated from the samples of soil seed banks, including in total 320 seedlings of *Sporobolus cryptandrus*. Beside of *Sporobolus*, *Arenaria leptoclados/serpyllifolia, Portulaca oleracea, Potentilla argentea, Digitaria sanguinalis* and *Cerastium semidecandrum* were the most frequent species in the seed bank with 508, 492, 200, 153 and 104 seedlings, respectively. These six taxa provided more than 83% of the total seed bank (see more details in Electronic Appendix 3). The seed density of *Sporobolus* ranged from 1,114 to 3,077 seeds/m^2^ considering the pooled seed bank data for the 30 cores (0-10 cm layer) per site (Electronic Appendix 3). Increasing *Sporobolus* cover negatively affected the total seedling number of other species, and also the species richness, Shannon diversity and evenness of the seed bank (Table 4, Figures 8, 9 and 10). The cover of *Sporobolus* did not significantly affect the seed bank density of *Sporobolus* itself; we detected soil seed banks of *Sporobolus* even in the closely located reference stands with no cover of the species. The soil layer significantly affected almost all seed bank characteristics, with decreasing scores towards to the deeper soil layers (Figures 8, 9 and 10). Seedlings of *Sporobolus* emerged from all studied soil layers (Figure 7). Only a few cases showed interaction between the soil layers and *Sporobolus* cover – in case of the seed bank density of *Sporobolus* and evenness (Table 4).

**Figure 7.**
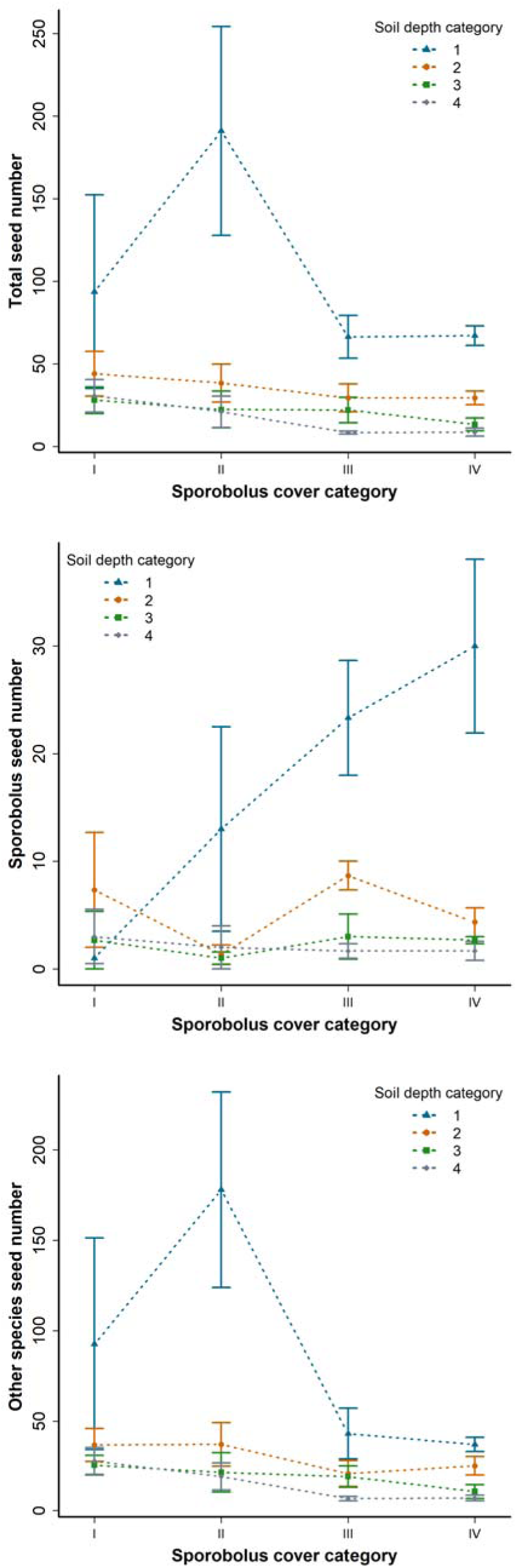
The relationship between *Sporobolus* cover categories and total seed bank density, density of the seed bank of *Sporobolus* and other species.

**Figure 8.**
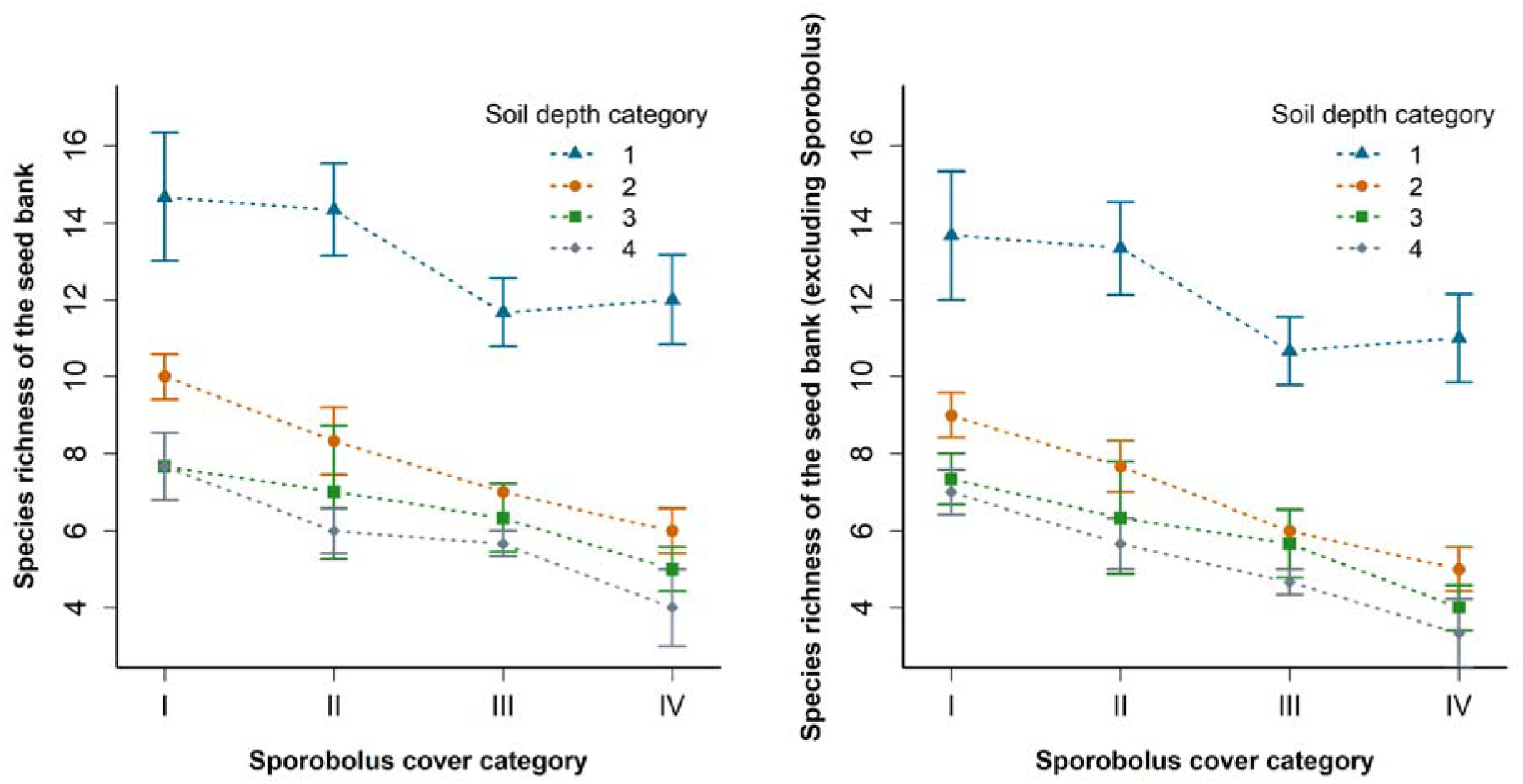
The relationship between *Sporobolus* cover categories and species richness in the seed banks.

**Figure 9.**
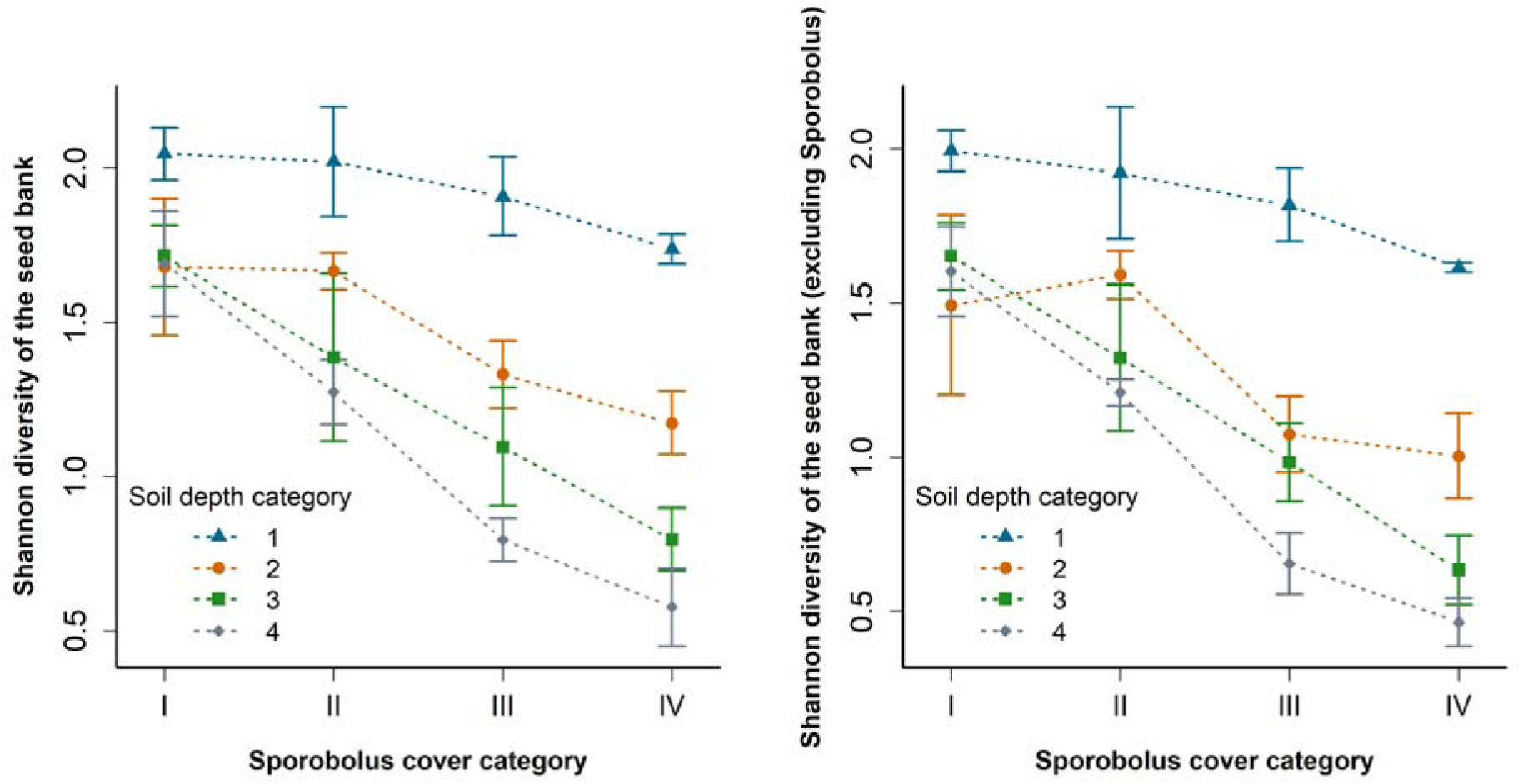
The relationship between *Sporobolus* cover categories and the Shannon diversity of the seed banks.

**Figure 10.**
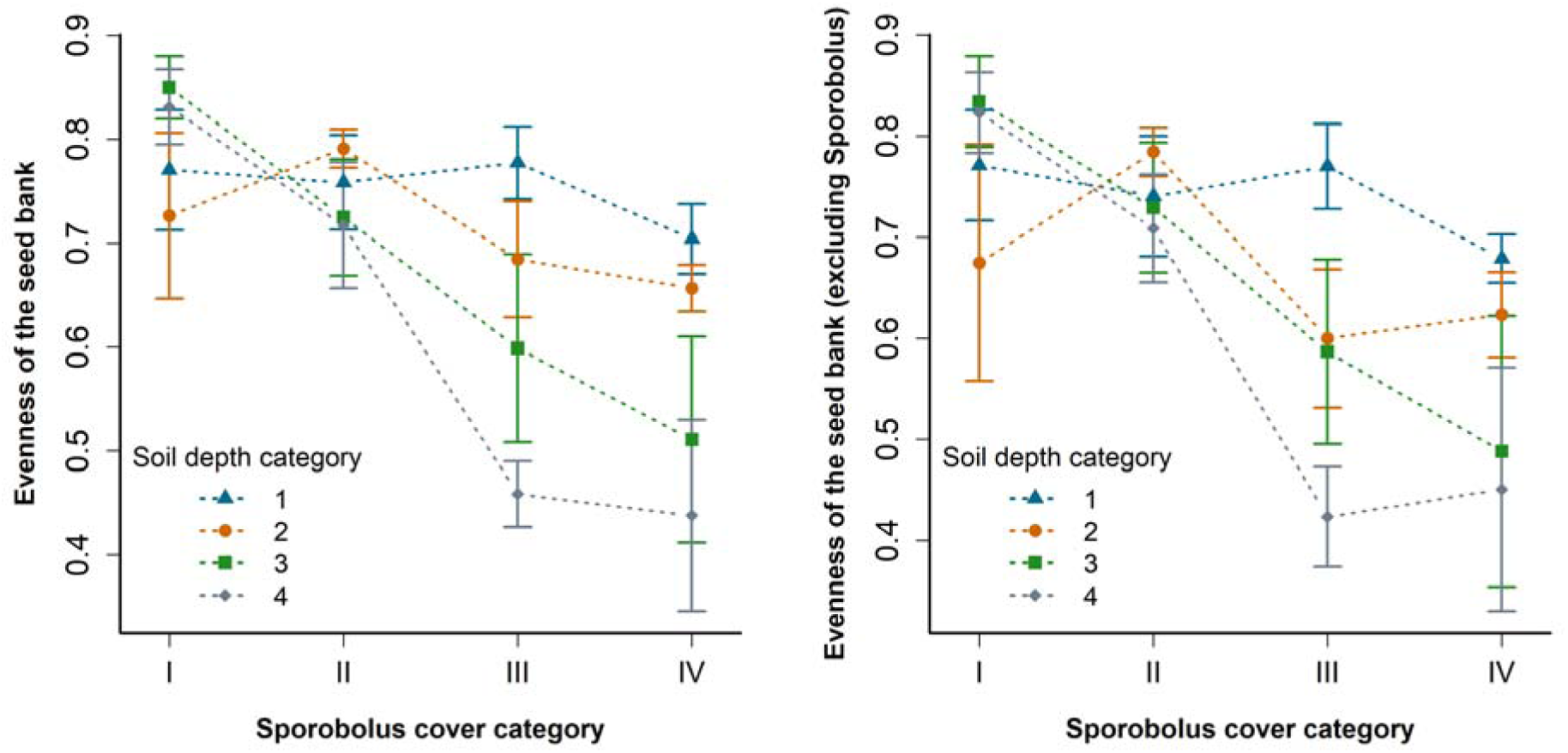
The relationship between *Sporobolus* cover categories and the evenness of the seed bank.

**Table 4.** Effect of *Sporobolus cryptandrus* cover on seed bank composition of the subjected plots. Significant effects were denoted with boldface (two-way ANOVA).

| Characteristic | *Sporobolus* cover |  | Soil layer |  | *Sporobolus* cover<br>× Soil layer |  |
| --- | --- | --- | --- | --- | --- | --- |
|  | *F*<sub>3,47</sub> | *p* | *F*<sub>3,47</sub> | *p* | *F*<sub>9,47</sub> | *p* |
| **Seedling number** |  |  |  |  |  |  |
| Total | 2.442 | 0.082 | **12.465** | **<0.001** | 1.503 | 0.189 |
| *Sporobolus* | 2.727 | 0.060 | **12.845** | **<0.001** | **2.913** | **0.012** |
| Other species | **3.848** | **0.019** | **10.133** | **<0.001** | 2.024 | 0.069 |
| **including *Sporobolus*** |  |  |  |  |  |  |
| Species richness | **8.697** | **<0.001** | **47.602** | **<0.001** | 0.312 | 0.965 |
| Shannon diversity | **19.698** | **<0.001** | **26.435** | **<0.001** | 1.419 | 0.222 |
| Evenness | **11.914** | **<0.001** | **4.320** | **0.011** | **2.277** | **0.042** |
| **excluding *Sporobolus*** |  |  |  |  |  |  |
| Species richness | **11.626** | **<0.001** | **50.652** | **<0.001** | 0.264 | 0.980 |
| Shannon diversity | **23.346** | **<0.001** | **26.845** | **<0.001** | 1.622 | 0.151 |
| Evenness | **8.659** | **<0.001** | 2.482 | 0.079 | 1.612 | 0.154 |

### Germination characteristics

Nearly 24% of all seeds germinated during the experiment. The highest total germination rate was detected in pots with low soil burial depth and no litter cover (Figure 11). Both litter and soil cover significantly affected the total germination rate and the seedling survival in the germination experiment, but there was no interaction between these factors (Table 5). The total germination rate was the lowest in pots with low to high burial depth and high litter cover, but even from these pots some seedlings appeared at the surface and were established until the end of the project. Considering all litter treatments, the highest total germination rate was detected with low soil cover and not in the case of no soil cover. Seedling survival rates showed high fluctuations in most treatments, even in treatments with relatively high survival rates, there were pots with relatively low survival rates. The lowest mean survival rates were detected for the treatments no soil/no litter and high soil/high litter cover (Figure 11).

**Figure 11.**
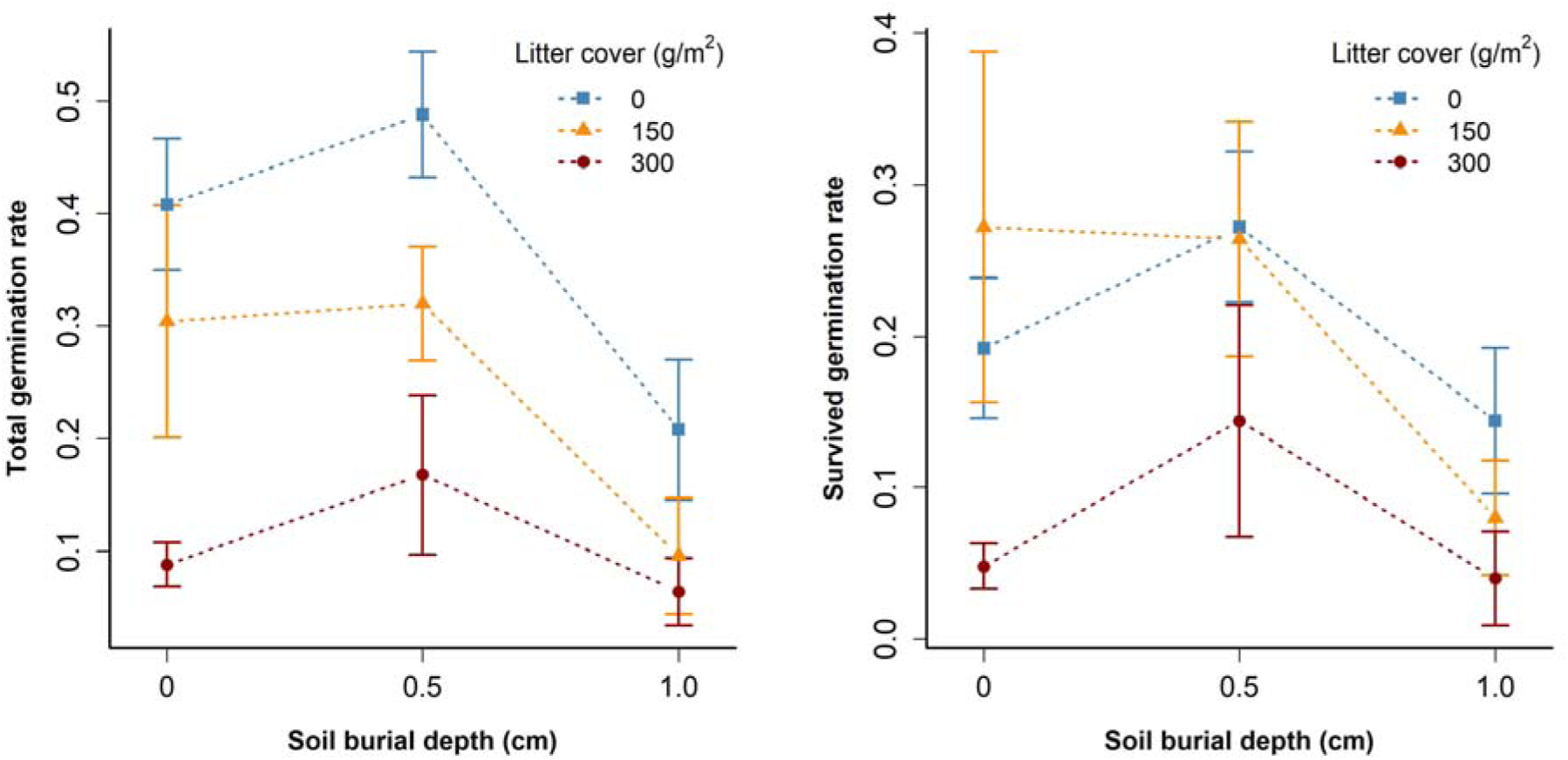
Effect of litter and soil covering on the total number of germinated seedlings (left) and seedling survival (right) of *Sporobolus cryptandrus*.

**Table 5.**
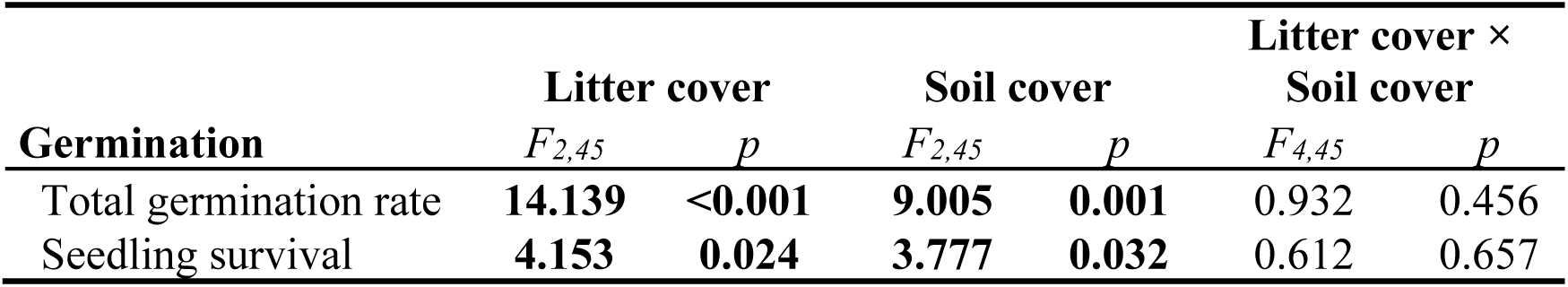
Effect of litter and soil cover on total germination rate and seedling survival of *Sporobolus cryptandrus* (two-way ANOVA, significant values are indicated with boldface, p<0.05).

| **Germination** | **Litter cover** |  | **Soil cover** |  | **Litter cover ×**<br>**Soil cover** |  |
| --- | --- | --- | --- | --- | --- | --- |
|  | *F*<sub>2,45</sub> | *p* | *F*<sub>2,45</sub> | *p* | *F*<sub>4,45</sub> | *p* |
| Total germination rate | **14.139** | **<0.001** | **9.005** | **0.001** | 0.932 | 0.456 |
| Seedling survival | **4.153** | **0.024** | **3.777** | **0.032** | 0.612 | 0.657 |

## Discussion

### Distribution and invasion of Sporobolus crypandrus

In recent decades, spread of several non-native *Sporobolus* species was detected in Europe. Beside *Sporobolus cryptandrus*, the establishment and intensive spread of naturalised populations of *S. neglectus* and *S. vaginiflorus* was reported from the Mediterranean regions and from dry regions of the eastern part of Central Europe (Hohla et al. 2015, Király & Hohla 2015, Jogan 2017, Englmaier & Wilhalm 2018). Rapid spread and expansion in the last decade were detected for both species along regularly mown margins of roads and motorways, but, because of the circumstances of their establishment, they constitute a lower threat to natural vegetation than *Sporobolus cryptandrus*. As in the case of *S. neglectus and S. vaginiflorus*, the current distribution map of *S. cryptandrus* clearly shows that the naturalised populations of this species are confined to the European Mediterranean or to regions of Eastern Central and Eastern Europe characterised by arid, at least moderately continental climate. Only occasional establishment of the species was detected in more humid regions of Central and Western Europe (Figure 2). In contrast to the other two species, spread of *S. cryptandrus* is not limited to road margins and the vicinity of ruderal sites (Hohla et al. 2015, but occasional establishment can occur also in isolated locations - see also Király 2016), but subjects large areas characterised by natural, semi-natural and degraded dry-grassland vegetation (this paper and Török & Aradi 2017). In the Kiskunság region, the species has also established in relatively undisturbed sandy grasslands. It is especially alarming that the species has established also in steppe grasslands in the Ukrainian and Russian steppe regions (Demina et al. 2018). As *S. cryptandrus* is considered to be one of the most drought-resistant species of short-grass prairies (see for example Tilley et al. 2009), further potential occurrences and its spread can be forecasted in dry sand regions or degraded rocky habitats of Europe due to the ongoing climate change.

### Effect of S. cryptandrus on the vegetation and seed banks of sand grasslands

Species-specific information on the aspects of the seed bank formation and early establishment patterns of an invasive species is crucial for developing strategies for its suppression and for the prevention of its further spread (Gioria et al. 2012, Sonkoly et al. 2020). We found that increasing cover of *S. cryptandrus* decreases the species richness and abundance of subordinate species both in the vegetation and seed banks. We also found a rather weak but facilitative effect of the low-abundance establishment of *S. cryptandrus* on the species richness of other species both in the vegetation and seed banks of the subjected grasslands. Similar facilitative effects were detected by Kelemen et al. (2015) in the case of the native species *Festuca pseudovina*, a community dominant perennial grass characteristic in dry alkali grasslands. The most likely explanation of the phenomenon could be that the establishment of the drought-tolerant species also mitigates the microclimatic extremities of the dry habitat and thus facilitate the establishment and survival of others (e.g., Eviner 2004) – especially that of short-lived species in the habitat (e.g., *Arenaria leptoclados*, *Portulaca oleracea* or *Cerastium semidecandrum*). However, this ‘nurse’ effect was not detected in plots with a high cover of the species. The facilitative effect of dominant perennials (including, in our case, *Sporobolus*) mostly occurs through the facilitation of the germination and early establishment of subordinate species, but this positive interaction can turn into competition for light or space (Liancourt et al. 2005, Le Roux et al. 2013). The sign of the interaction between plant species is also density-dependent; a low density of a facilitator species can have positive effects, but it can turn to negative interaction above a certain density of the species (Kelemen et al. 2019). This is in line with our findings as the richness and abundance of subordinated species in the vegetation and seed banks was higher in plots with a low density of *Sporobolus* than in plots without *Sporobolus,* but a higher density of the species had an overall negative effect.

Viable seeds of the species were detected from all soil layers. This indicates that the species is able to form a persistent seed bank as also indicated by similar results from natural prairie communities in North America (Coffin & Lauenroth 1989, Pérez et al. 1998, Clements et al. 2007). However, the seed bank density was rather low in the three deeper soil layers and increasing *Sporobolus* abundance in the vegetation only increased *Sporobolus* seed bank density in the uppermost layer. This may be an indirect evidence of the ongoing spread and quite recent establishment of the species in the study site (i.e., only a small number of seeds were able to reach the deeper soil layers in the limited time). The detected seed density scores for *Sporobolus* are comparable to some studies in native prairie habitats where high densities of the species were typical (up to 3,414 seeds/m^2^, Clements et al. 2007). It cannot be excluded, however, that the frequent mowing at the sampled urban grassland occurring before seed maturation affects the seed production and seed bank accumulation of the species in spite of its ability of late and secondary flowering. This latter issue should be clarified later when the seed banks of all sites will be assessed.

### Effect of litter and soil cover on the germination of the species

We found that germination of *Sporobolus cryptandrus* was negatively affected by soil burial and litter cover, but there was no interaction between the two factors. Some seedlings emerged in pots even with the highest litter and soil cover levels. These results were in line with most findings of Sonkoly et al. (2020) where the germination of 11 invasive species were tested. Mirroring also the results of the latter study, the small-seeded *S. cryptandrus* had the highest seed germination rates with 0.5 cm soil burial depth with no litter cover. The detected seed germination rates of *Sporobolus* were rather low compared to some other invasive grasses, but quite similar and even a bit higher compared to the formerly reported germination/viability rates after warm stratification (Sartor & Malone 2010). As seeds were collected in previous years, these results suggest that the seeds can survive even longer periods of dry storage without the significant loss of their viability.

### Open research questions and conservation outlook

By definition, those species can be considered as transformer invasive species “that change the character, condition, form, or nature of ecosystems” (Richardson et al. 2000). Based on our results, *Sporobolus cryptandrus* can be considered as a transformer invasive species, whose spread poses a high risk for dry sand and steppe grasslands in Eurasia, especially in the steppe climatic zone. However, to develop an appropriate strategy for its suppression we need further information on the following crucial aspects of its life history, population dynamics and spread. First, in depth genetic analyses (e.g., a phylogeographic study using a genomic method) are needed to clarify the likely origin of the established populations and to interpret the means of its long-distance dispersal making it possible to evaluate the pace of its spread and population growth. Second, we need further information on its competitive ability including not only data on the aboveground competition but also information of its allelopathic ability and root competition. Third, it would be necessary to assess the seed banks of sites with different site history and establishment time of the species to evaluate the density and the development speed of its soil seed banks. Lastly, for its effective suppression it is vital to study its possible enemies in its native area and to assess the capacity of traditional management types (grazing or mowing) to control the species, not excluding the possibility of its eradication using mechanical (e.g., shading) and/or chemical methods.

## Acknowledgements

PT, ZB and CT were supported by NKFIH (K119225, K124796 and PD132131, respectively) and the ‘Momentum’ Program of the Hungarian Academy of Sciences during manuscript writing.

---

**Electronic Appendix 1.**
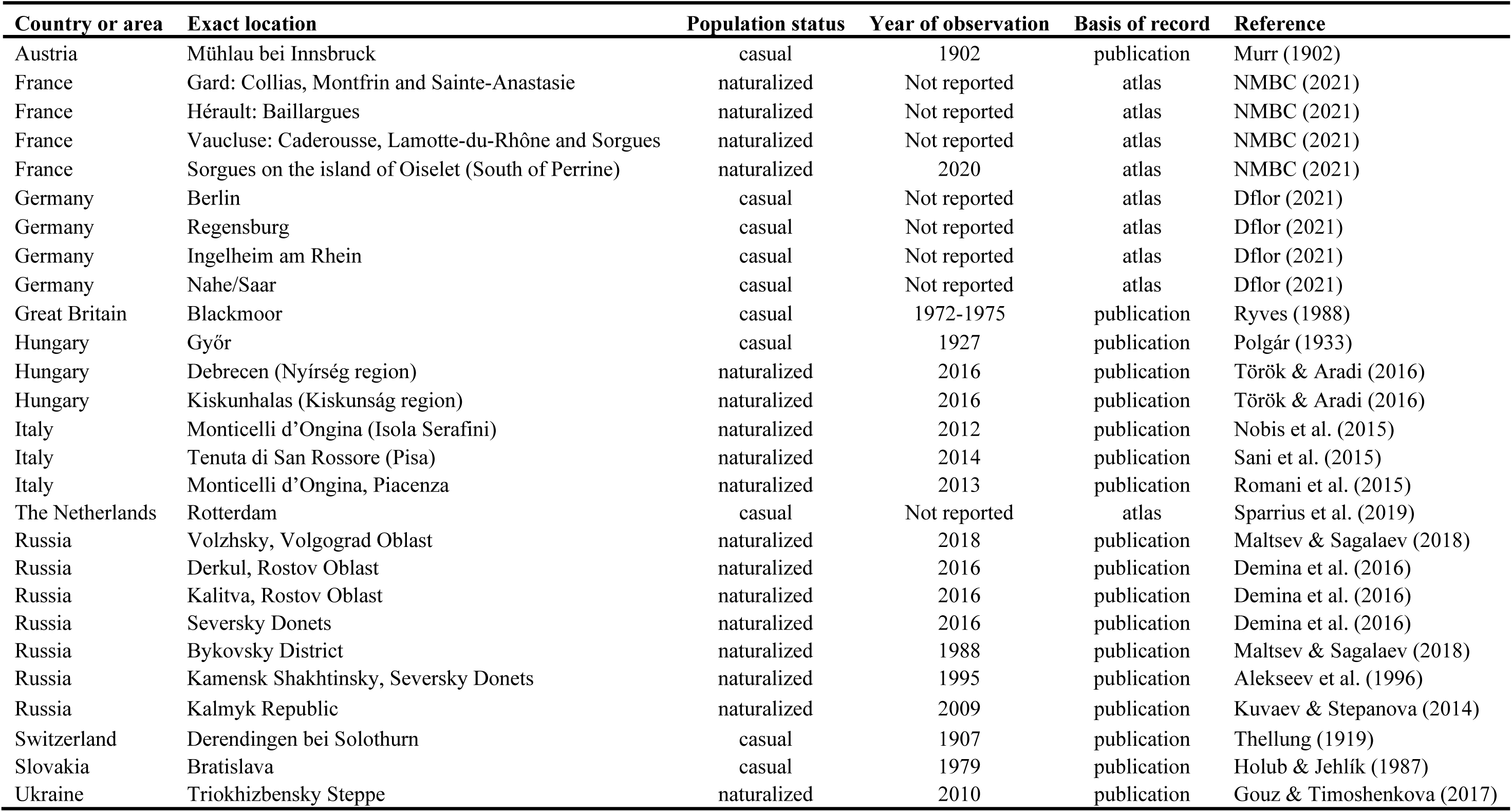
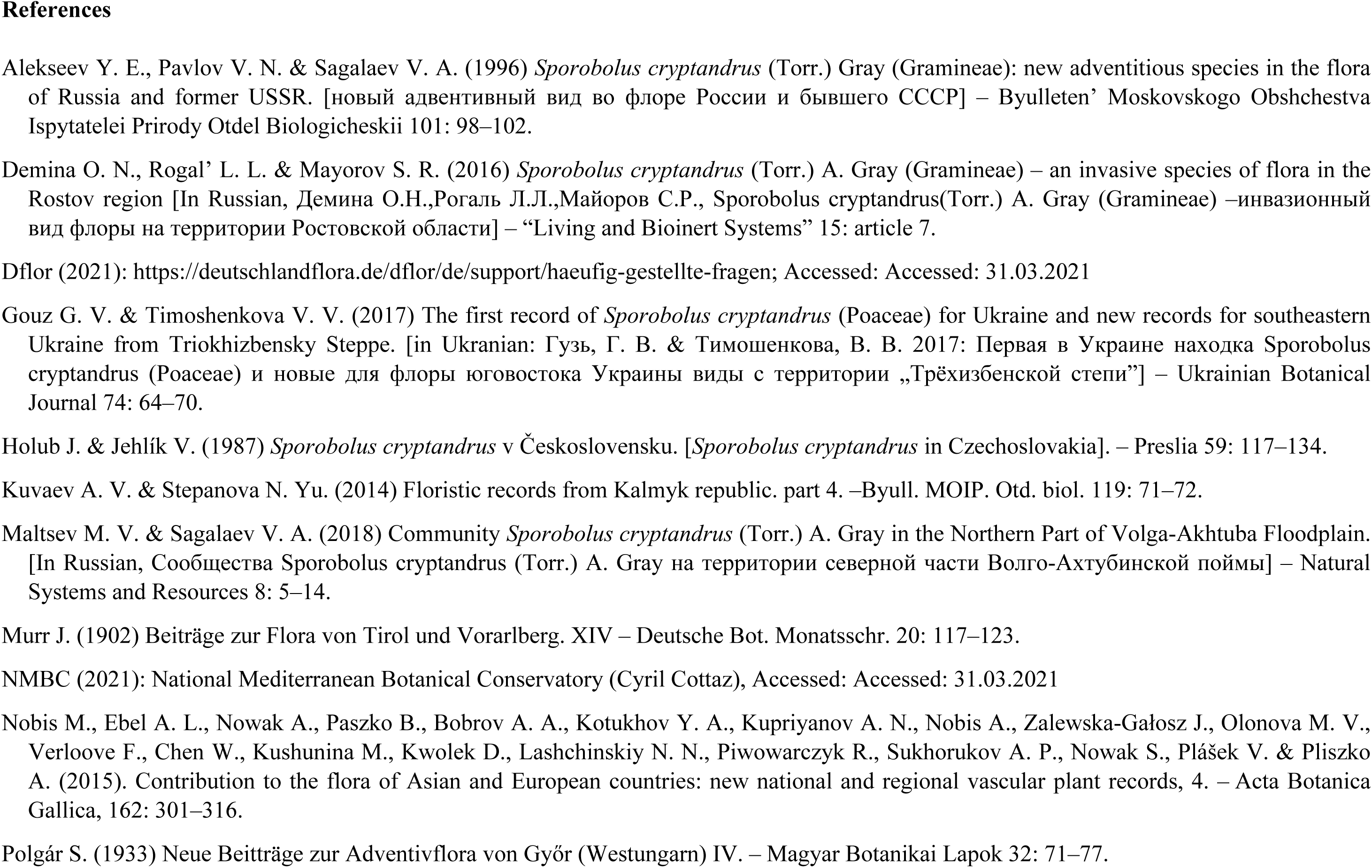
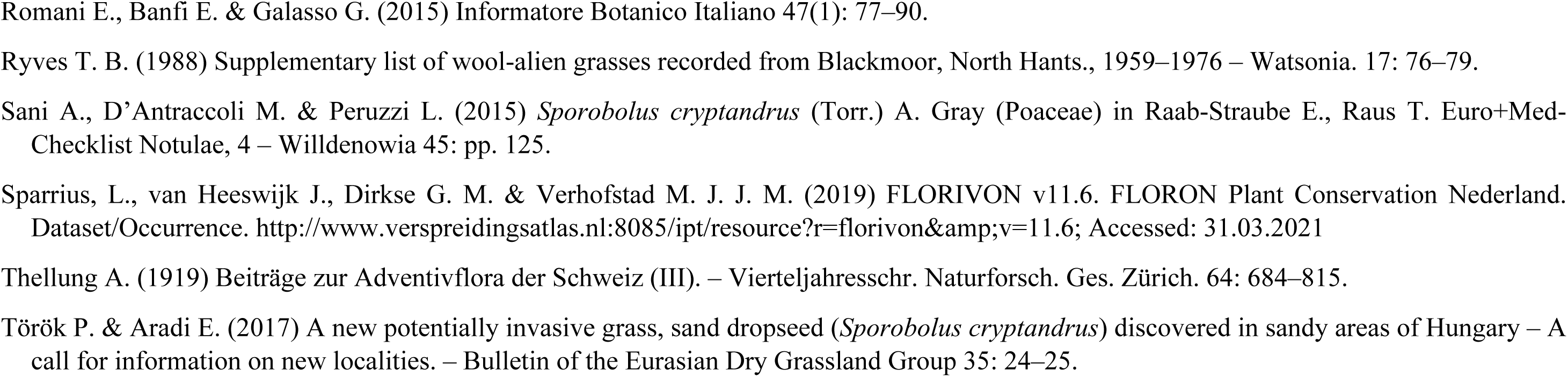
The distribution of *Sporobolus cryptandrus* in Eurasia based on published literature data. For the locations visualised on a map please see also Figure 2.

**Electronic Appendix 2A.**
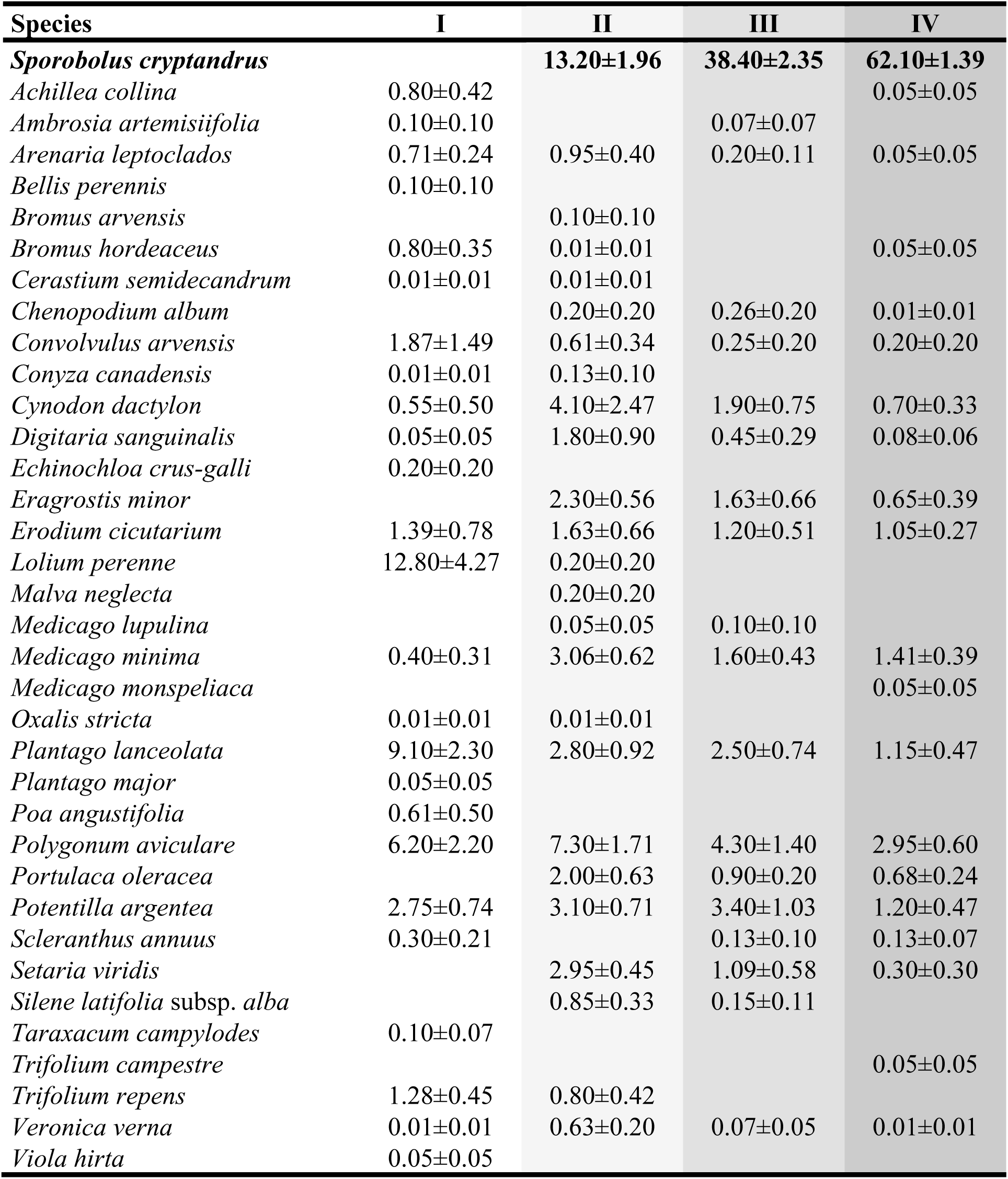
Species composition of the vegetation at the Debrecen study site. Cover categories: I: nearby reference sites where *Sporobolus* is missing, II: 1-25% of *Sporobolus* cover, III: 26-50% of *Sporobolus* cover, IV >50% of *Sporobolus* cover. In each cover category the cover scores of species in 10, 1-m^2^-sized plots were averaged (mean±SE).

**Electronic Appendix 2B.**
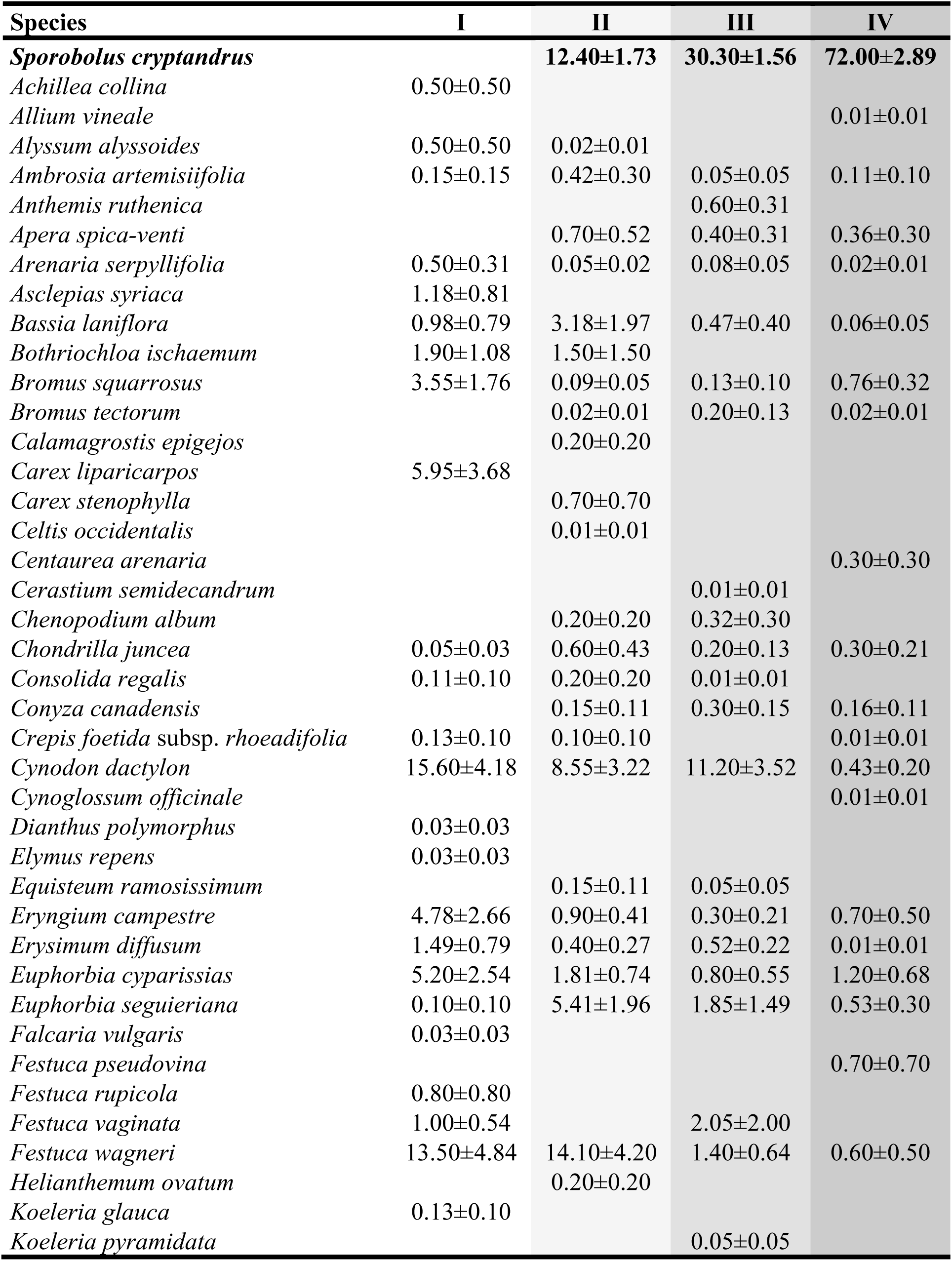
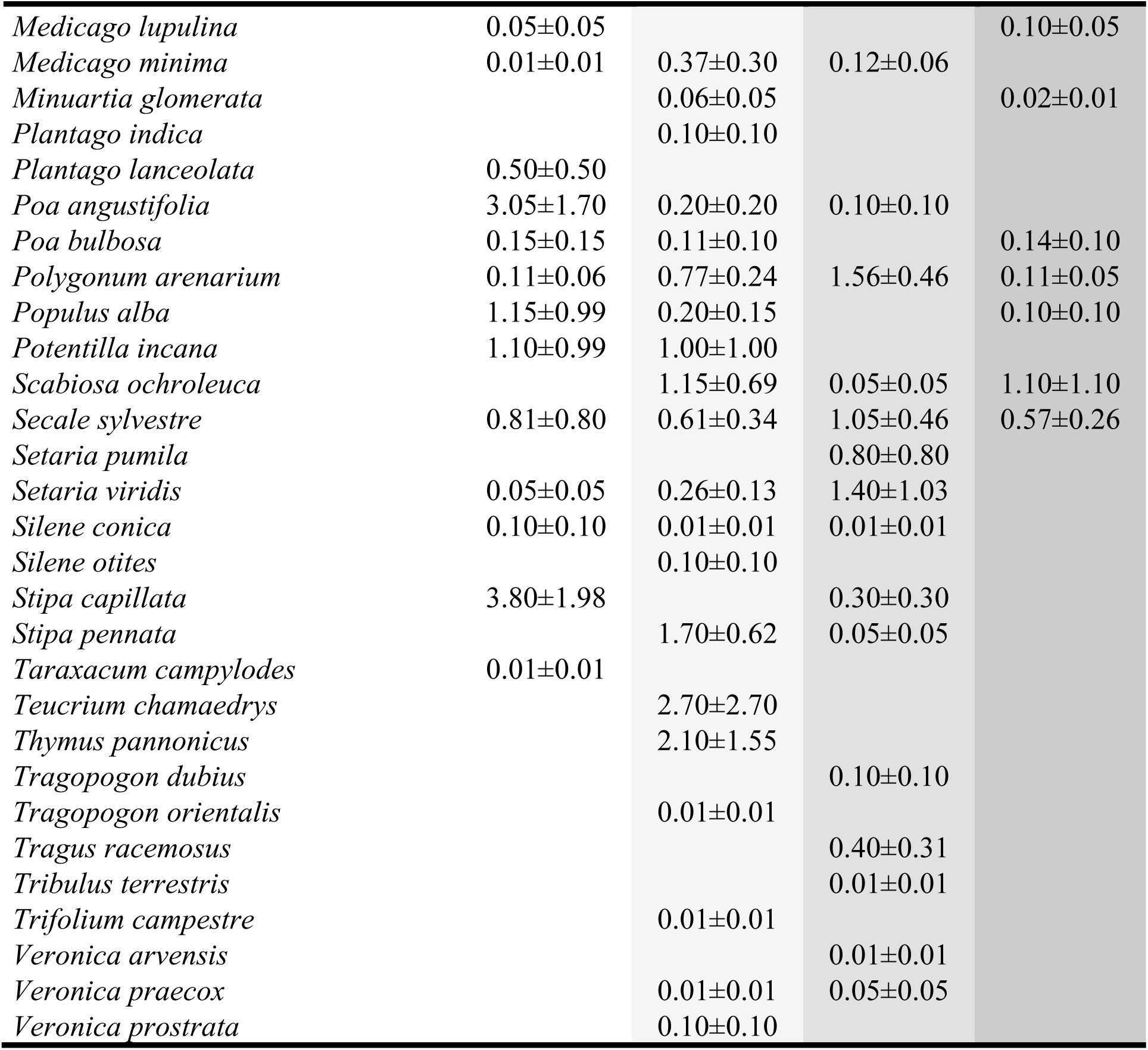
Species composition of vegetation at the Kiskunhalas North (KN) study site. Cover categories: I: nearby reference sites where *Sporobolus* is missing, II: 1-25% of *Sporobolus* cover, III: 26-50% of *Sporobolus* cover, IV >50% of *Sporobolus* cover. In each cover category the cover scores of species in 10, 1-m^2^-sized plots were averaged (mean±SE).

**Electronic Appendix 2C.**
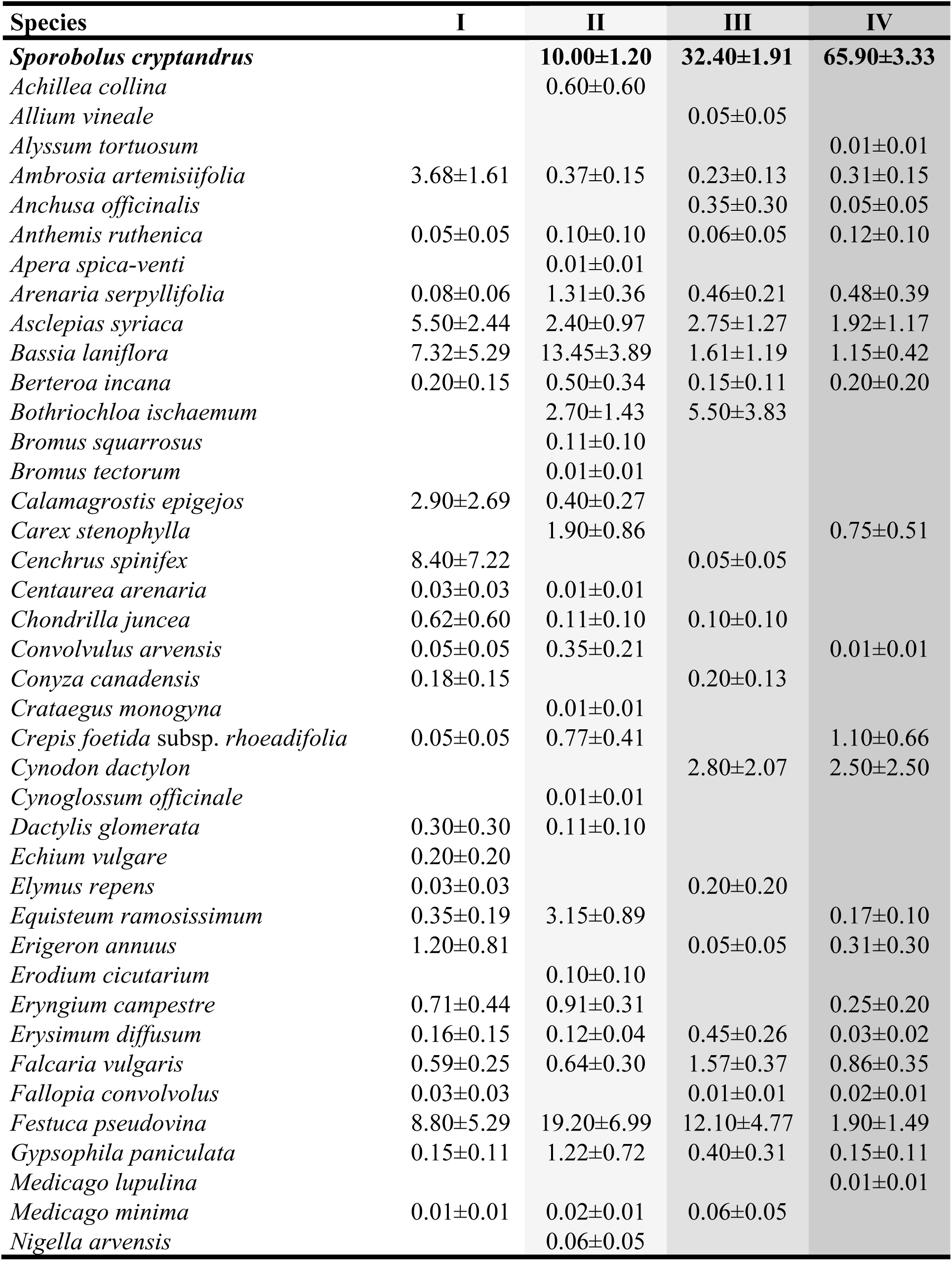
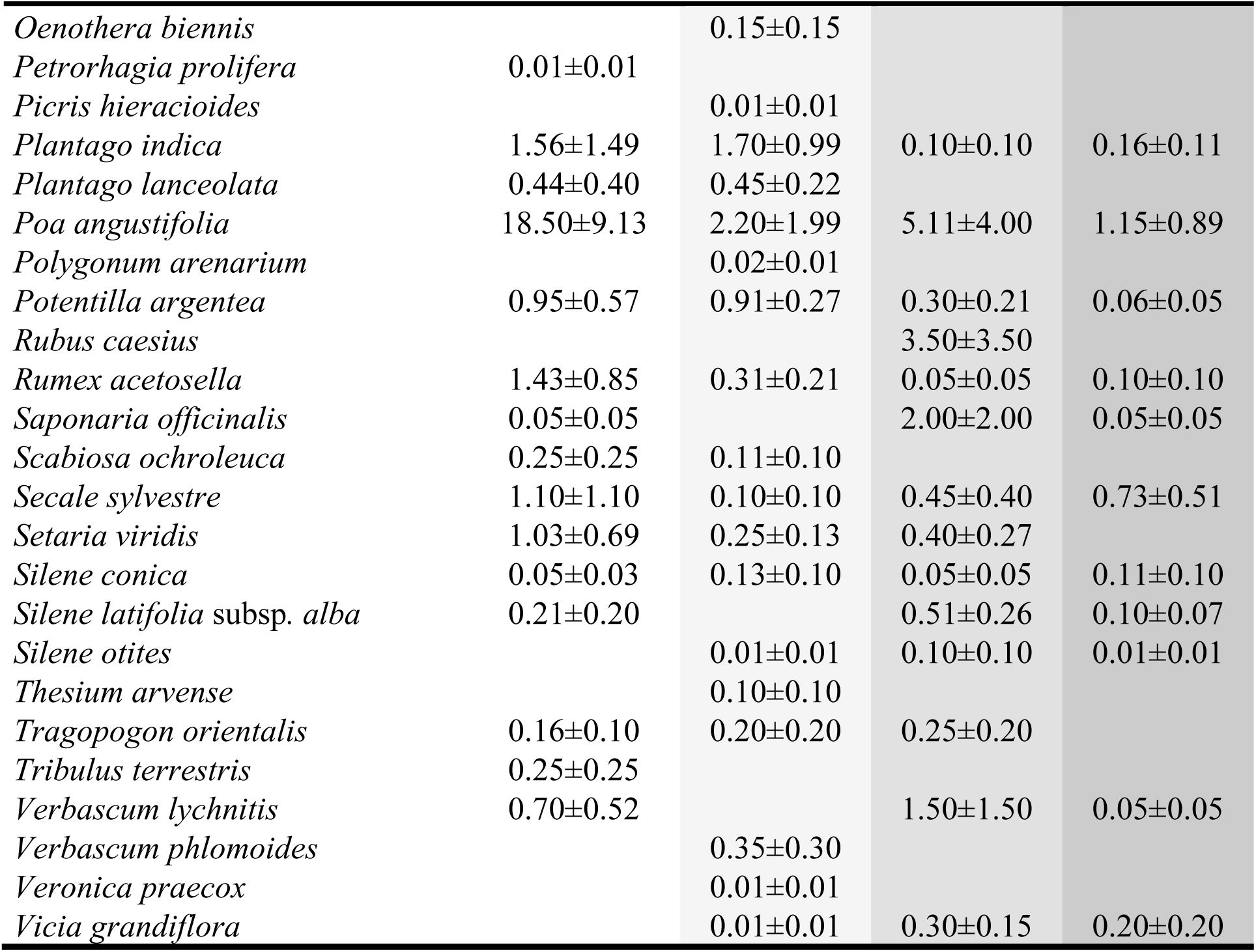
Species composition of vegetation at the Katonatelep (KT) study site. Cover categories: I: nearby reference sites where *Sporobolus* is missing, II: 1-25% of *Sporobolus* cover, III: 26-50% of *Sporobolus* cover, IV >50% of *Sporobolus* cover. In each cover category the cover scores of species of 10, 1-m^2^-sized plots were averaged (mean±SE).

**Electronic Appendix 2D.**
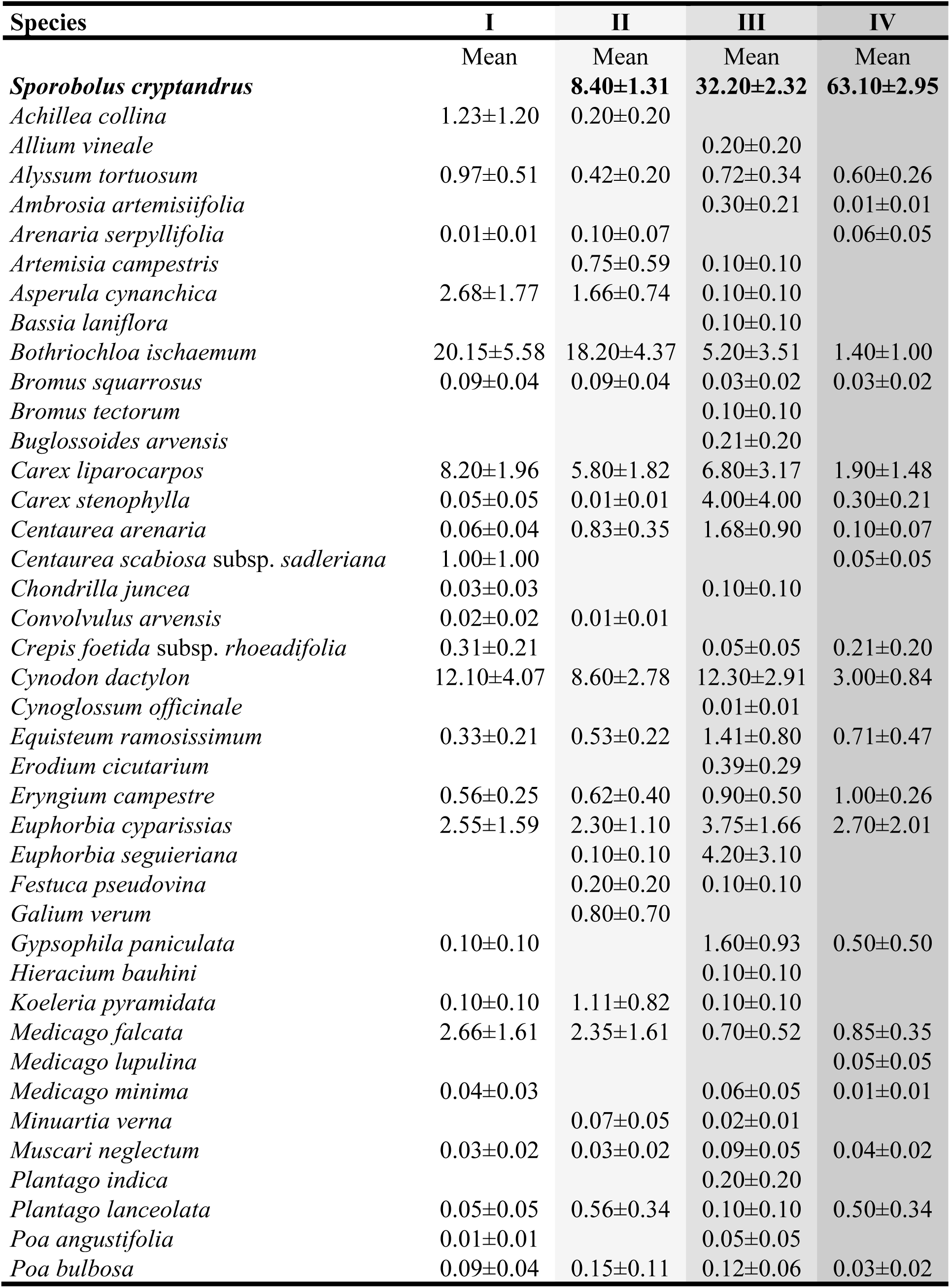
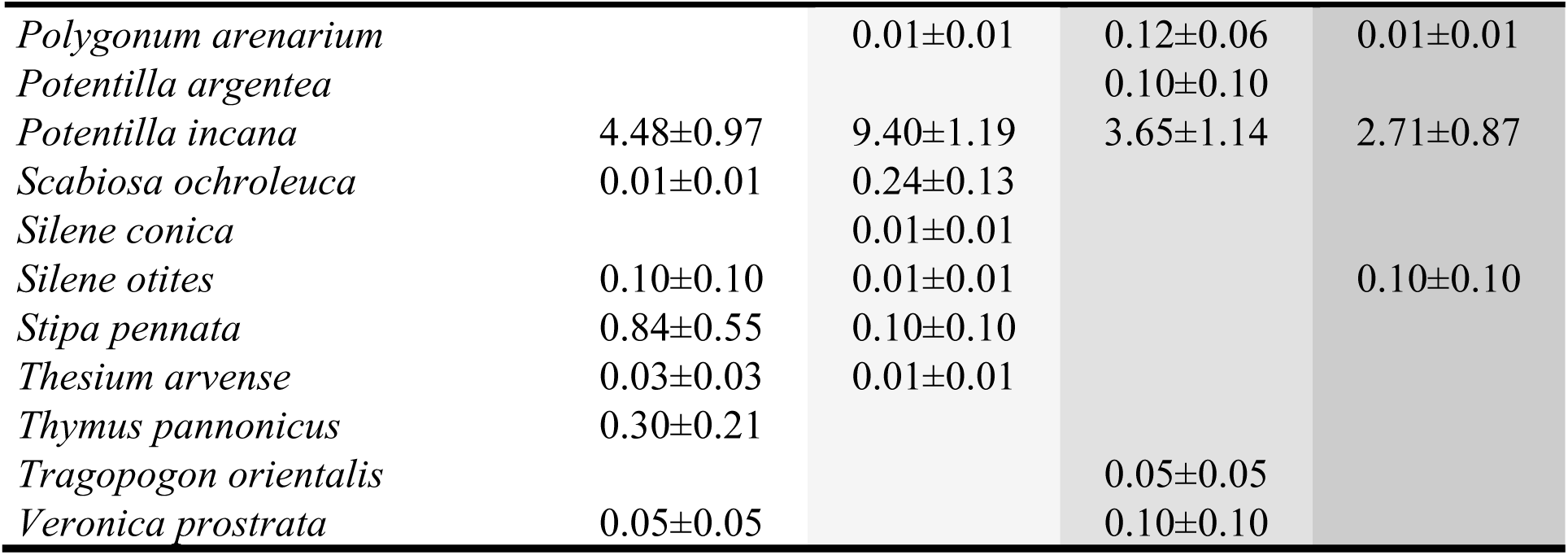
Species composition of the vegetation at the Airport (A) study site. Cover categories: I: nearby reference sites where *Sporobolus* is missing, II: 1-25% of *Sporobolus* cover, III: 26-50% of *Sporobolus* cover, IV >50% of *Sporobolus* cover. In each cover category the cover scores of species of 10, 1-m^2^-sized plots were averaged (mean±SE).

**Electronic Appendix 2E.**
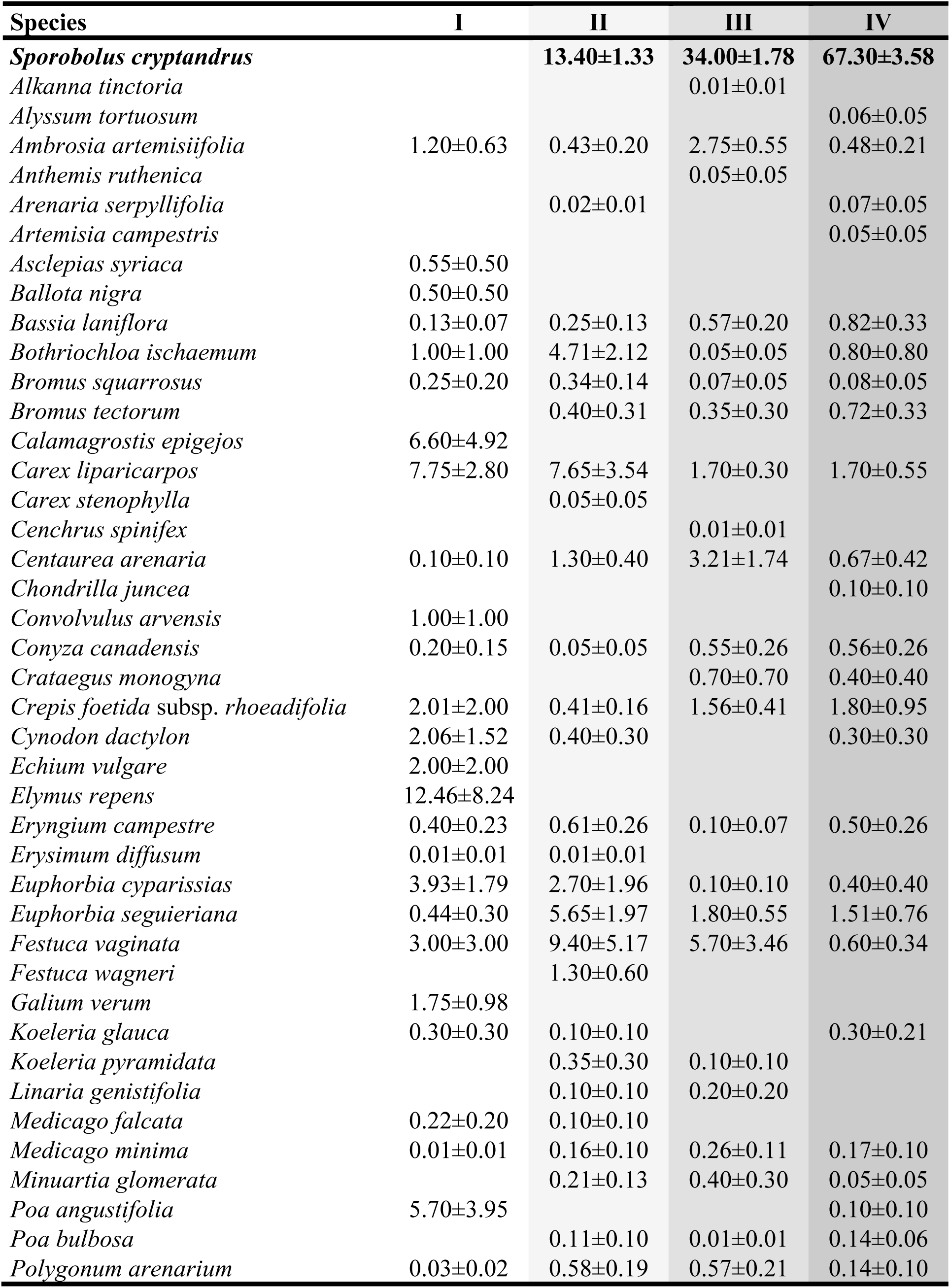
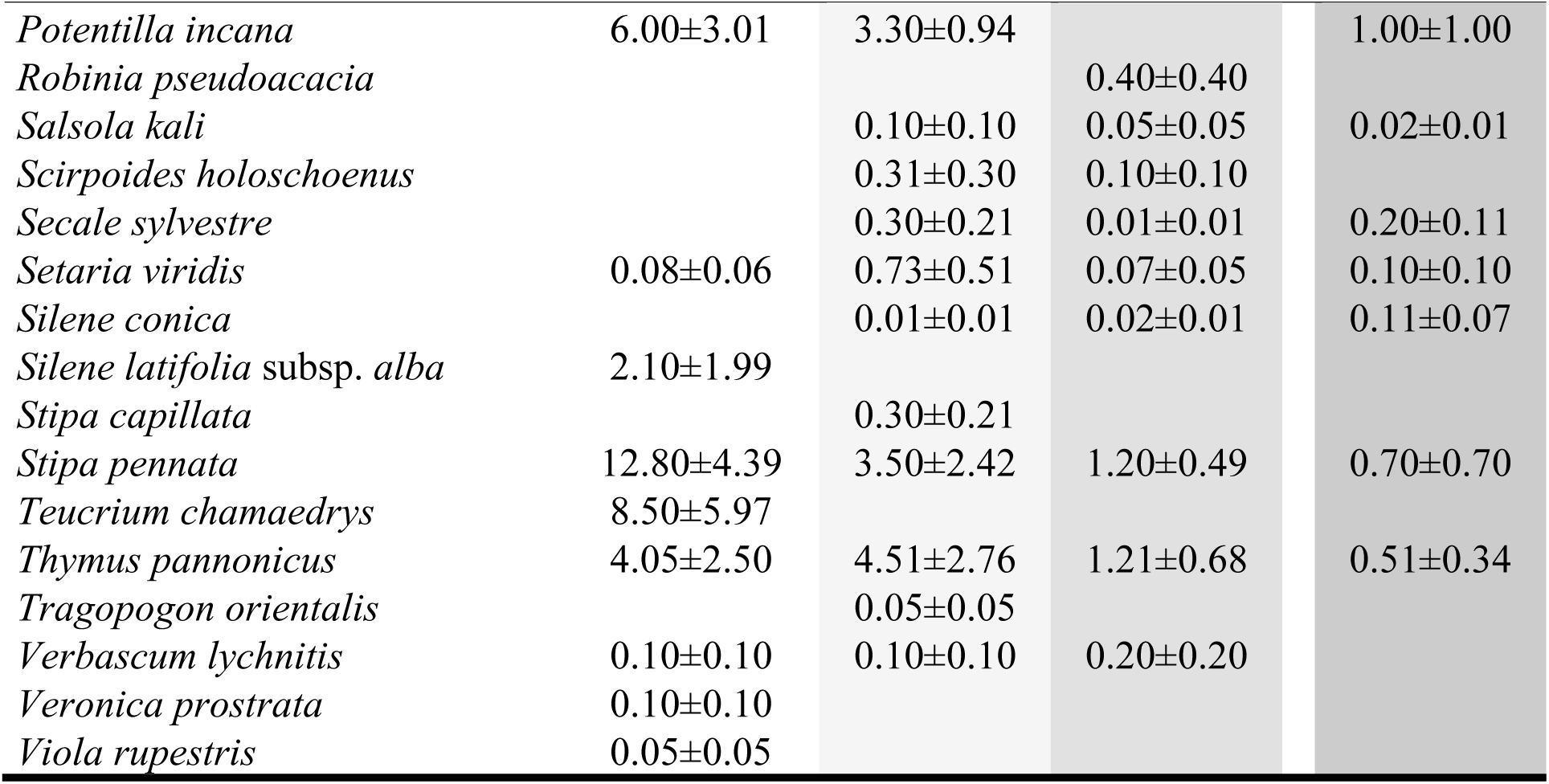
Species composition of vegetation at the Kiskunhalas East study site. Cover categories: I: nearby reference sites where *Sporobolus* is missing, II: 1-25% of *Sporobolus* cover, III: 26-50% of *Sporobolus* cover, IV >50% of *Sporobolus* cover. In each cover category the cover scores of species of 10, 1-m^2^-sized plots were averaged (mean±SE).

**Electronic Appendix 3.**
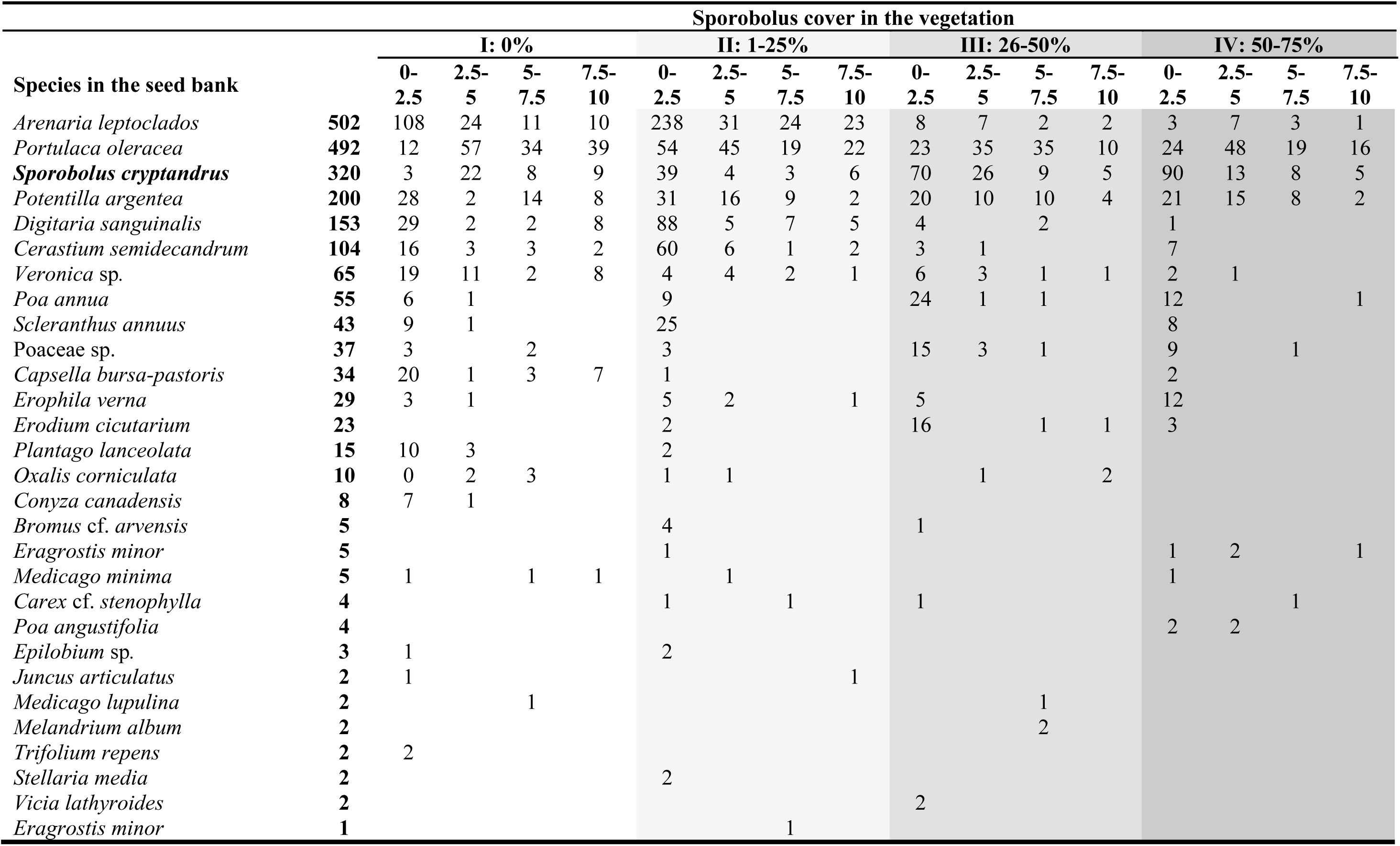
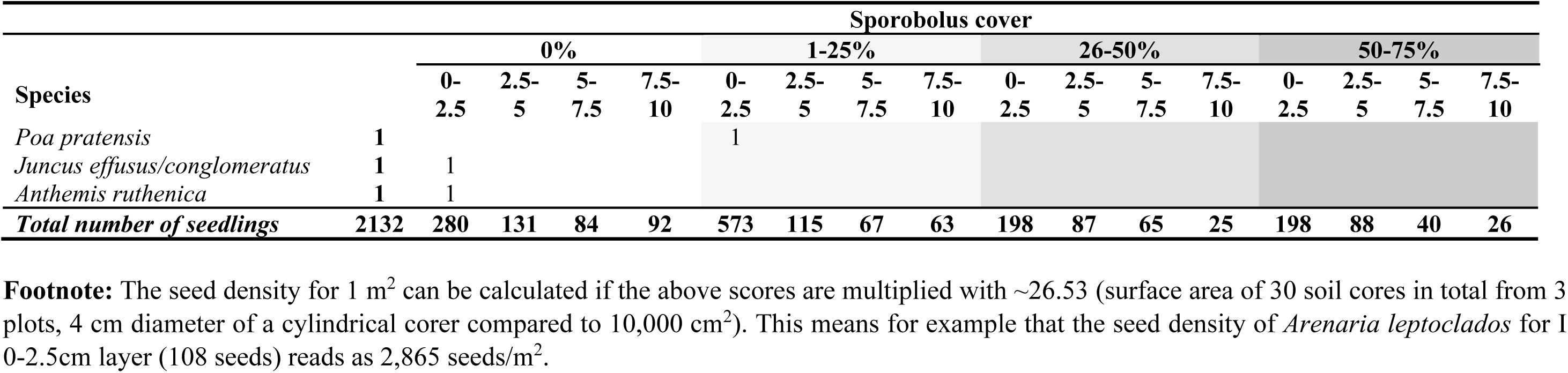
Seed bank composition of the sites with an increasing cover of *Sporobolus cryptandrus*. In the table seedling numbers are shown.

