## Supplementary material for "Sand dropseed (*Sporobolus cryptandrus*) – A new pest in Eurasian sand areas?": E-Appendix1

**Electronic Appendix 1.** The distribution of *Sporobolus cryptandrus* in Eurasia based on published literature data. For the locations visualised on a map please see also Figure 2.

| Country or area | Exact location | Population status | Year of observation | Basis of record | Reference |
|---|---|---|---|---|---|
| Austria | Mühlau bei Innsbruck | casual | 1902 | publication | Murr (1902) |
| France | Gard: Collias, Montfrin and Sainte-Anastasie | naturalized | Not reported | atlas | NMBC (2021) |
| France | Hérault: Baillargues | naturalized | Not reported | atlas | NMBC (2021) |
| France | Vaucluse: Caderousse, Lamotte-du-Rhône and Sorgues | naturalized | Not reported | atlas | NMBC (2021) |
| France | Sorgues on the island of Oiselet (South of Perrine) | naturalized | 2020 | atlas | NMBC (2021) |
| Germany | Berlin | casual | Not reported | atlas | Dflor (2021) |
| Germany | Regensburg | casual | Not reported | atlas | Dflor (2021) |
| Germany | Ingelheim am Rhein | casual | Not reported | atlas | Dflor (2021) |
| Germany | Nahe/Saar | casual | Not reported | atlas | Dflor (2021) |
| Great Britain | Blackmoor | casual | 1972-1975 | publication | Ryves (1988) |
| Hungary | Győr | casual | 1927 | publication | Polgár (1933) |
| Hungary | Debrecen (Nyírség region) | naturalized | 2016 | publication | Török & Aradi (2016) |
| Hungary | Kiskunhalas (Kiskunság region) | naturalized | 2016 | publication | Török & Aradi (2016) |
| Italy | Monticelli d'Ongina (Isola Serafini) | naturalized | 2012 | publication | Nobis et al. (2015) |
| Italy | Tenuta di San Rossore (Pisa) | naturalized | 2014 | publication | Sani et al. (2015) |
| Italy | Monticelli d'Ongina, Piacenza | naturalized | 2013 | publication | Romani et al. (2015) |
| The Netherlands | Rotterdam | casual | Not reported | atlas | Sparrius et al. (2019) |
| Russia | Volzhsky, Volgograd Oblast | naturalized | 2018 | publication | Maltsev & Sagalaev (2018) |
| Russia | Derkul, Rostov Oblast | naturalized | 2016 | publication | Demina et al. (2016) |
| Russia | Kalitva, Rostov Oblast | naturalized | 2016 | publication | Demina et al. (2016) |
| Russia | Seversky Donets | naturalized | 2016 | publication | Demina et al. (2016) |
| Russia | Bykovsky District | naturalized | 1988 | publication | Maltsev & Sagalaev (2018) |
| Russia | Kamensk Shakhtinsky, Seversky Donets | naturalized | 1995 | publication | Alekseev et al. (1996) |
| Russia | Kalmyk Republic | naturalized | 2009 | publication | Kuvaev & Stepanova (2014) |
| Switzerland | Derendingen bei Solothurn | casual | 1907 | publication | Thellung (1919) |
| Slovakia | Bratislava | casual | 1979 | publication | Holub & Jehlík (1987) |
| Ukraine | Triokhizbensky Steppe | naturalized | 2010 | publication | Gouz & Timoshenkova (2017) |
