## Supplementary material for "Sand dropseed (*Sporobolus cryptandrus*) – A new pest in Eurasian sand areas?": E-Appendix2

**Electronic Appendix 2A.** Species composition of the vegetation at the Debrecen study site. Cover categories: I: nearby reference sites where *Sporobolus* is missing, II: 1-25% of *Sporobolus* cover, III: 26-50% of *Sporobolus* cover, IV >50% of *Sporobolus* cover. In each cover category the cover scores of species in 10, 1-$m^2$-sized plots were averaged (mean±SE).

| **Species** | **I** | **II** | **III** | **IV** |
|---|---|---|---|---|
| ***Sporobolus cryptandrus*** | | **13.20±1.96** | **38.40±2.35** | **62.10±1.39** |
| *Achillea collina* | 0.80±0.42 | | | 0.05±0.05 |
| *Ambrosia artemisiifolia* | 0.10±0.10 | | 0.07±0.07 | |
| *Arenaria leptoclados* | 0.71±0.24 | 0.95±0.40 | 0.20±0.11 | 0.05±0.05 |
| *Bellis perennis* | 0.10±0.10 | | | |
| *Bromus arvensis* | | 0.10±0.10 | | |
| *Bromus hordeaceus* | 0.80±0.35 | 0.01±0.01 | | 0.05±0.05 |
| *Cerastium semidecandrum* | 0.01±0.01 | 0.01±0.01 | | |
| *Chenopodium album* | | 0.20±0.20 | 0.26±0.20 | 0.01±0.01 |
| *Convolvulus arvensis* | 1.87±1.49 | 0.61±0.34 | 0.25±0.20 | 0.20±0.20 |
| *Conyza canadensis* | 0.01±0.01 | 0.13±0.10 | | |
| *Cynodon dactylon* | 0.55±0.50 | 4.10±2.47 | 1.90±0.75 | 0.70±0.33 |
| *Digitaria sanguinalis* | 0.05±0.05 | 1.80±0.90 | 0.45±0.29 | 0.08±0.06 |
| *Echinochloa crus-galli* | 0.20±0.20 | | | |
| *Eragrostis minor* | | 2.30±0.56 | 1.63±0.66 | 0.65±0.39 |
| *Erodium cicutarium* | 1.39±0.78 | 1.63±0.66 | 1.20±0.51 | 1.05±0.27 |
| *Lolium perenne* | 12.80±4.27 | 0.20±0.20 | | |
| *Malva neglecta* | | 0.20±0.20 | | |
| *Medicago lupulina* | | 0.05±0.05 | 0.10±0.10 | |
| *Medicago minima* | 0.40±0.31 | 3.06±0.62 | 1.60±0.43 | 1.41±0.39 |
| *Medicago monspeliaca* | | | | 0.05±0.05 |
| *Oxalis stricta* | 0.01±0.01 | 0.01±0.01 | | |
| *Plantago lanceolata* | 9.10±2.30 | 2.80±0.92 | 2.50±0.74 | 1.15±0.47 |
| *Plantago major* | 0.05±0.05 | | | |
| *Poa angustifolia* | 0.61±0.50 | | | |
| *Polygonum aviculare* | 6.20±2.20 | 7.30±1.71 | 4.30±1.40 | 2.95±0.60 |
| *Portulaca oleracea* | | 2.00±0.63 | 0.90±0.20 | 0.68±0.24 |
| *Potentilla argentea* | 2.75±0.74 | 3.10±0.71 | 3.40±1.03 | 1.20±0.47 |
| *Scleranthus annuus* | 0.30±0.21 | | 0.13±0.10 | 0.13±0.07 |
| *Setaria viridis* | | 2.95±0.45 | 1.09±0.58 | 0.30±0.30 |
| *Silene latifolia* subsp. *alba* | | 0.85±0.33 | 0.15±0.11 | |
| *Taraxacum campylodes* | 0.10±0.07 | | | |
| *Trifolium campestre* | | | | 0.05±0.05 |
| *Trifolium repens* | 1.28±0.45 | 0.80±0.42 | | |
| *Veronica verna* | 0.01±0.01 | 0.63±0.20 | 0.07±0.05 | 0.01±0.01 |
| *Viola hirta* | 0.05±0.05 | | | |

**Electronic Appendix 2B.** Species composition of vegetation at the Kiskunhalas North (KN) study site. Cover categories: I: nearby reference sites where *Sporobolus* is missing, II: 1-25% of *Sporobolus* cover, III: 26-50% of *Sporobolus* cover, IV >50% of *Sporobolus* cover. In each cover category the cover scores of species in 10, 1-m$^2$-sized plots were averaged (mean±SE).

| **Species** | **I** | **II** | **III** | **IV** |
|---|---|---|---|---|
| ***Sporobolus cryptandrus*** | | **12.40±1.73** | **30.30±1.56** | **72.00±2.89** |
| *Achillea collina* | 0.50±0.50 | | | |
| *Allium vineale* | | | | 0.01±0.01 |
| *Alyssum alyssoides* | 0.50±0.50 | 0.02±0.01 | | |
| *Ambrosia artemisiifolia* | 0.15±0.15 | 0.42±0.30 | 0.05±0.05 | 0.11±0.10 |
| *Anthemis ruthenica* | | | 0.60±0.31 | |
| *Apera spica-venti* | | 0.70±0.52 | 0.40±0.31 | 0.36±0.30 |
| *Arenaria serpyllifolia* | 0.50±0.31 | 0.05±0.02 | 0.08±0.05 | 0.02±0.01 |
| *Asclepias syriaca* | 1.18±0.81 | | | |
| *Bassia laniflora* | 0.98±0.79 | 3.18±1.97 | 0.47±0.40 | 0.06±0.05 |
| *Bothriochloa ischaemum* | 1.90±1.08 | 1.50±1.50 | | |
| *Bromus squarrosus* | 3.55±1.76 | 0.09±0.05 | 0.13±0.10 | 0.76±0.32 |
| *Bromus tectorum* | | 0.02±0.01 | 0.20±0.13 | 0.02±0.01 |
| *Calamagrostis epigejos* | | 0.20±0.20 | | |
| *Carex liparicarpos* | 5.95±3.68 | | | |
| *Carex stenophylla* | | 0.70±0.70 | | |
| *Celtis occidentalis* | | 0.01±0.01 | | |
| *Centaurea arenaria* | | | | 0.30±0.30 |
| *Cerastium semidecandrum* | | | 0.01±0.01 | |
| *Chenopodium album* | | 0.20±0.20 | 0.32±0.30 | |
| *Chondrilla juncea* | 0.05±0.03 | 0.60±0.43 | 0.20±0.13 | 0.30±0.21 |
| *Consolida regalis* | 0.11±0.10 | 0.20±0.20 | 0.01±0.01 | |
| *Conyza canadensis* | | 0.15±0.11 | 0.30±0.15 | 0.16±0.11 |
| *Crepis foetida* subsp. *rhoeadifolia* | 0.13±0.10 | 0.10±0.10 | | 0.01±0.01 |
| *Cynodon dactylon* | 15.60±4.18 | 8.55±3.22 | 11.20±3.52 | 0.43±0.20 |
| *Cynoglossum officinale* | | | | 0.01±0.01 |
| *Dianthus polymorphus* | 0.03±0.03 | | | |
| *Elymus repens* | 0.03±0.03 | | | |
| *Equisteum ramosissimum* | | 0.15±0.11 | 0.05±0.05 | |
| *Eryngium campestre* | 4.78±2.66 | 0.90±0.41 | 0.30±0.21 | 0.70±0.50 |
| *Erysimum diffusum* | 1.49±0.79 | 0.40±0.27 | 0.52±0.22 | 0.01±0.01 |
| *Euphorbia cyparissias* | 5.20±2.54 | 1.81±0.74 | 0.80±0.55 | 1.20±0.68 |
| *Euphorbia seguieriana* | 0.10±0.10 | 5.41±1.96 | 1.85±1.49 | 0.53±0.30 |
| *Falcaria vulgaris* | 0.03±0.03 | | | |
| *Festuca pseudovina* | | | | 0.70±0.70 |
| *Festuca rupicola* | 0.80±0.80 | | | |
| *Festuca vaginata* | 1.00±0.54 | | 2.05±2.00 | |
| *Festuca wagneri* | 13.50±4.84 | 14.10±4.20 | 1.40±0.64 | 0.60±0.50 |
| *Helianthemum ovatum* | | 0.20±0.20 | | |
| *Koeleria glauca* | 0.13±0.10 | | | |
| *Koeleria pyramidata* | | | 0.05±0.05 | |

**Electronic Appendix 2B. continued.**

| Species | I | II | III | IV |
|---|---|---|---|---|
| *Medicago lupulina* | 0.05±0.05 | | | 0.10±0.05 |
| *Medicago minima* | 0.01±0.01 | 0.37±0.30 | 0.12±0.06 | |
| *Minuartia glomerata* | | 0.06±0.05 | | 0.02±0.01 |
| *Plantago indica* | | 0.10±0.10 | | |
| *Plantago lanceolata* | 0.50±0.50 | | | |
| *Poa angustifolia* | 3.05±1.70 | 0.20±0.20 | 0.10±0.10 | |
| *Poa bulbosa* | 0.15±0.15 | 0.11±0.10 | | 0.14±0.10 |
| *Polygonum arenarium* | 0.11±0.06 | 0.77±0.24 | 1.56±0.46 | 0.11±0.05 |
| *Populus alba* | 1.15±0.99 | 0.20±0.15 | | 0.10±0.10 |
| *Potentilla incana* | 1.10±0.99 | 1.00±1.00 | | |
| *Scabiosa ochroleuca* | | 1.15±0.69 | 0.05±0.05 | 1.10±1.10 |
| *Secale sylvestre* | 0.81±0.80 | 0.61±0.34 | 1.05±0.46 | 0.57±0.26 |
| *Setaria pumila* | | | 0.80±0.80 | |
| *Setaria viridis* | 0.05±0.05 | 0.26±0.13 | 1.40±1.03 | |
| *Silene conica* | 0.10±0.10 | 0.01±0.01 | 0.01±0.01 | |
| *Silene otites* | | 0.10±0.10 | | |
| *Stipa capillata* | 3.80±1.98 | | 0.30±0.30 | |
| *Stipa pennata* | | 1.70±0.62 | 0.05±0.05 | |
| *Taraxacum campylodes* | 0.01±0.01 | | | |
| *Teucrium chamaedrys* | | 2.70±2.70 | | |
| *Thymus pannonicus* | | 2.10±1.55 | | |
| *Tragopogon dubius* | | | 0.10±0.10 | |
| *Tragopogon orientalis* | | 0.01±0.01 | | |
| *Tragus racemosus* | | | 0.40±0.31 | |
| *Tribulus terrestris* | | | 0.01±0.01 | |
| *Trifolium campestre* | | 0.01±0.01 | | |
| *Veronica arvensis* | | | 0.01±0.01 | |
| *Veronica praecox* | | 0.01±0.01 | 0.05±0.05 | |
| *Veronica prostrata* | | 0.10±0.10 | | |

**Electronic Appendix 2C.** Species composition of vegetation at the Katonatelep (KT) study site. Cover categories: I: nearby reference sites where *Sporobolus* is missing, II: 1-25% of *Sporobolus* cover, III: 26-50% of *Sporobolus* cover, IV >50% of *Sporobolus* cover. In each cover category the cover scores of species of 10, 1-$m^2$-sized plots were averaged (mean±SE).

| **Species** | **I** | **II** | **III** | **IV** |
|---|---|---|---|---|
| ***Sporobolus cryptandrus*** | | **10.00±1.20** | **32.40±1.91** | **65.90±3.33** |
| *Achillea collina* | | 0.60±0.60 | | |
| *Allium vineale* | | | 0.05±0.05 | |
| *Alyssum tortuosum* | | | | 0.01±0.01 |
| *Ambrosia artemisiifolia* | 3.68±1.61 | 0.37±0.15 | 0.23±0.13 | 0.31±0.15 |
| *Anchusa officinalis* | | | 0.35±0.30 | 0.05±0.05 |
| *Anthemis ruthenica* | 0.05±0.05 | 0.10±0.10 | 0.06±0.05 | 0.12±0.10 |
| *Apera spica-venti* | | 0.01±0.01 | | |
| *Arenaria serpyllifolia* | 0.08±0.06 | 1.31±0.36 | 0.46±0.21 | 0.48±0.39 |
| *Asclepias syriaca* | 5.50±2.44 | 2.40±0.97 | 2.75±1.27 | 1.92±1.17 |
| *Bassia laniflora* | 7.32±5.29 | 13.45±3.89 | 1.61±1.19 | 1.15±0.42 |
| *Berteroa incana* | 0.20±0.15 | 0.50±0.34 | 0.15±0.11 | 0.20±0.20 |
| *Bothriochloa ischaemum* | | 2.70±1.43 | 5.50±3.83 | |
| *Bromus squarrosus* | | 0.11±0.10 | | |
| *Bromus tectorum* | | 0.01±0.01 | | |
| *Calamagrostis epigejos* | 2.90±2.69 | 0.40±0.27 | | |
| *Carex stenophylla* | | 1.90±0.86 | | 0.75±0.51 |
| *Cenchrus spinifex* | 8.40±7.22 | | 0.05±0.05 | |
| *Centaurea arenaria* | 0.03±0.03 | 0.01±0.01 | | |
| *Chondrilla juncea* | 0.62±0.60 | 0.11±0.10 | 0.10±0.10 | |
| *Convolvulus arvensis* | 0.05±0.05 | 0.35±0.21 | | 0.01±0.01 |
| *Conyza canadensis* | 0.18±0.15 | | 0.20±0.13 | |
| *Crataegus monogyna* | | 0.01±0.01 | | |
| *Crepis foetida* subsp. *rhoeadifolia* | 0.05±0.05 | 0.77±0.41 | | 1.10±0.66 |
| *Cynodon dactylon* | | | 2.80±2.07 | 2.50±2.50 |
| *Cynoglossum officinale* | | 0.01±0.01 | | |
| *Dactylis glomerata* | 0.30±0.30 | 0.11±0.10 | | |
| *Echium vulgare* | 0.20±0.20 | | | |
| *Elymus repens* | 0.03±0.03 | | 0.20±0.20 | |
| *Equisteum ramosissimum* | 0.35±0.19 | 3.15±0.89 | | 0.17±0.10 |
| *Erigeron annuus* | 1.20±0.81 | | 0.05±0.05 | 0.31±0.30 |
| *Erodium cicutarium* | | 0.10±0.10 | | |
| *Eryngium campestre* | 0.71±0.44 | 0.91±0.31 | | 0.25±0.20 |
| *Erysimum diffusum* | 0.16±0.15 | 0.12±0.04 | 0.45±0.26 | 0.03±0.02 |
| *Falcaria vulgaris* | 0.59±0.25 | 0.64±0.30 | 1.57±0.37 | 0.86±0.35 |
| *Fallopia convolvolus* | 0.03±0.03 | | 0.01±0.01 | 0.02±0.01 |
| *Festuca pseudovina* | 8.80±5.29 | 19.20±6.99 | 12.10±4.77 | 1.90±1.49 |
| *Gypsophila paniculata* | 0.15±0.11 | 1.22±0.72 | 0.40±0.31 | 0.15±0.11 |
| *Medicago lupulina* | | | | 0.01±0.01 |
| *Medicago minima* | 0.01±0.01 | 0.02±0.01 | 0.06±0.05 | |
| *Nigella arvensis* | | 0.06±0.05 | | |

**Electronic Appendix 2C continued.**

| **Species** | **I** | **II** | **III** | **IV** |
|---|---|---|---|---|
| *Oenothera biennis* | | 0.15±0.15 | | |
| *Petrorhagia prolifera* | 0.01±0.01 | | | |
| *Picris hieracioides* | | 0.01±0.01 | | |
| *Plantago indica* | 1.56±1.49 | 1.70±0.99 | 0.10±0.10 | 0.16±0.11 |
| *Plantago lanceolata* | 0.44±0.40 | 0.45±0.22 | | |
| *Poa angustifolia* | 18.50±9.13 | 2.20±1.99 | 5.11±4.00 | 1.15±0.89 |
| *Polygonum arenarium* | | 0.02±0.01 | | |
| *Potentilla argentea* | 0.95±0.57 | 0.91±0.27 | 0.30±0.21 | 0.06±0.05 |
| *Rubus caesius* | | | 3.50±3.50 | |
| *Rumex acetosella* | 1.43±0.85 | 0.31±0.21 | 0.05±0.05 | 0.10±0.10 |
| *Saponaria officinalis* | 0.05±0.05 | | 2.00±2.00 | 0.05±0.05 |
| *Scabiosa ochroleuca* | 0.25±0.25 | 0.11±0.10 | | |
| *Secale sylvestre* | 1.10±1.10 | 0.10±0.10 | 0.45±0.40 | 0.73±0.51 |
| *Setaria viridis* | 1.03±0.69 | 0.25±0.13 | 0.40±0.27 | |
| *Silene conica* | 0.05±0.03 | 0.13±0.10 | 0.05±0.05 | 0.11±0.10 |
| *Silene latifolia* subsp. *alba* | 0.21±0.20 | | 0.51±0.26 | 0.10±0.07 |
| *Silene otites* | | 0.01±0.01 | 0.10±0.10 | 0.01±0.01 |
| *Thesium arvense* | | 0.10±0.10 | | |
| *Tragopogon orientalis* | 0.16±0.10 | 0.20±0.20 | 0.25±0.20 | |
| *Tribulus terrestris* | 0.25±0.25 | | | |
| *Verbascum lychnitis* | 0.70±0.52 | | 1.50±1.50 | 0.05±0.05 |
| *Verbascum phlomoides* | | 0.35±0.30 | | |
| *Veronica praecox* | | 0.01±0.01 | | |
| *Vicia grandiflora* | | 0.01±0.01 | 0.30±0.15 | 0.20±0.20 |

**Electronic Appendix 2D.** Species composition of the vegetation at the Airport (A) study site. Cover categories: I: nearby reference sites where *Sporobolus* is missing, II: 1-25% of *Sporobolus* cover, III: 26-50% of *Sporobolus* cover, IV >50% of *Sporobolus* cover. In each cover category the cover scores of species of 10, 1-m$^2$-sized plots were averaged (mean±SE).

| **Species** | **I** | **II** | **III** | **IV** |
|---|---|---|---|---|
| | Mean | Mean | Mean | Mean |
| ***Sporobolus cryptandrus*** | | **8.40±1.31** | **32.20±2.32** | **63.10±2.95** |
| *Achillea collina* | 1.23±1.20 | 0.20±0.20 | | |
| *Allium vineale* | | | 0.20±0.20 | |
| *Alyssum tortuosum* | 0.97±0.51 | 0.42±0.20 | 0.72±0.34 | 0.60±0.26 |
| *Ambrosia artemisiifolia* | | | 0.30±0.21 | 0.01±0.01 |
| *Arenaria serpyllifolia* | 0.01±0.01 | 0.10±0.07 | | 0.06±0.05 |
| *Artemisia campestris* | | 0.75±0.59 | 0.10±0.10 | |
| *Asperula cynanchica* | 2.68±1.77 | 1.66±0.74 | 0.10±0.10 | |
| *Bassia laniflora* | | | 0.10±0.10 | |
| *Bothriochloa ischaemum* | 20.15±5.58 | 18.20±4.37 | 5.20±3.51 | 1.40±1.00 |
| *Bromus squarrosus* | 0.09±0.04 | 0.09±0.04 | 0.03±0.02 | 0.03±0.02 |
| *Bromus tectorum* | | | 0.10±0.10 | |
| *Buglossoides arvensis* | | | 0.21±0.20 | |
| *Carex liparocarpos* | 8.20±1.96 | 5.80±1.82 | 6.80±3.17 | 1.90±1.48 |
| *Carex stenophylla* | 0.05±0.05 | 0.01±0.01 | 4.00±4.00 | 0.30±0.21 |
| *Centaurea arenaria* | 0.06±0.04 | 0.83±0.35 | 1.68±0.90 | 0.10±0.07 |
| *Centaurea scabiosa* subsp. *sadleriana* | 1.00±1.00 | | | 0.05±0.05 |
| *Chondrilla juncea* | 0.03±0.03 | | 0.10±0.10 | |
| *Convolvulus arvensis* | 0.02±0.02 | 0.01±0.01 | | |
| *Crepis foetida* subsp. *rhoeadifolia* | 0.31±0.21 | | 0.05±0.05 | 0.21±0.20 |
| *Cynodon dactylon* | 12.10±4.07 | 8.60±2.78 | 12.30±2.91 | 3.00±0.84 |
| *Cynoglossum officinale* | | | 0.01±0.01 | |
| *Equisteum ramosissimum* | 0.33±0.21 | 0.53±0.22 | 1.41±0.80 | 0.71±0.47 |
| *Erodium cicutarium* | | | 0.39±0.29 | |
| *Eryngium campestre* | 0.56±0.25 | 0.62±0.40 | 0.90±0.50 | 1.00±0.26 |
| *Euphorbia cyparissias* | 2.55±1.59 | 2.30±1.10 | 3.75±1.66 | 2.70±2.01 |
| *Euphorbia seguieriana* | | 0.10±0.10 | 4.20±3.10 | |
| *Festuca pseudovina* | | 0.20±0.20 | 0.10±0.10 | |
| *Galium verum* | | 0.80±0.70 | | |
| *Gypsophila paniculata* | 0.10±0.10 | | 1.60±0.93 | 0.50±0.50 |
| *Hieracium bauhini* | | | 0.10±0.10 | |
| *Koeleria pyramidata* | 0.10±0.10 | 1.11±0.82 | 0.10±0.10 | |
| *Medicago falcata* | 2.66±1.61 | 2.35±1.61 | 0.70±0.52 | 0.85±0.35 |
| *Medicago lupulina* | | | | 0.05±0.05 |
| *Medicago minima* | 0.04±0.03 | | 0.06±0.05 | 0.01±0.01 |
| *Minuartia verna* | | 0.07±0.05 | 0.02±0.01 | |
| *Muscari neglectum* | 0.03±0.02 | 0.03±0.02 | 0.09±0.05 | 0.04±0.02 |
| *Plantago indica* | | | 0.20±0.20 | |
| *Plantago lanceolata* | 0.05±0.05 | 0.56±0.34 | 0.10±0.10 | 0.50±0.34 |
| *Poa angustifolia* | 0.01±0.01 | | 0.05±0.05 | |
| *Poa bulbosa* | 0.09±0.04 | 0.15±0.11 | 0.12±0.06 | 0.03±0.02 |

**Electronic Appendix 2D continued.**

| Species | I | II | III | IV |
|---|---|---|---|---|
| *Polygonum arenarium* | | 0.01±0.01 | 0.12±0.06 | 0.01±0.01 |
| *Potentilla argentea* | | | 0.10±0.10 | |
| *Potentilla incana* | 4.48±0.97 | 9.40±1.19 | 3.65±1.14 | 2.71±0.87 |
| *Scabiosa ochroleuca* | 0.01±0.01 | 0.24±0.13 | | |
| *Silene conica* | | 0.01±0.01 | | |
| *Silene otites* | 0.10±0.10 | 0.01±0.01 | | 0.10±0.10 |
| *Stipa pennata* | 0.84±0.55 | 0.10±0.10 | | |
| *Thesium arvense* | 0.03±0.03 | 0.01±0.01 | | |
| *Thymus pannonicus* | 0.30±0.21 | | | |
| *Tragopogon orientalis* | | | 0.05±0.05 | |
| *Veronica prostrata* | 0.05±0.05 | | 0.10±0.10 | |

**Electronic Appendix 2E.** Species composition of vegetation at the Kiskunhalas East study site. Cover categories: I: nearby reference sites where *Sporobolus* is missing, II: 1-25% of *Sporobolus* cover, III: 26-50% of *Sporobolus* cover, IV >50% of *Sporobolus* cover. In each cover category the cover scores of species of 10, 1-m$^2$-sized plots were averaged (mean±SE).

| **Species** | **I** | **II** | **III** | **IV** |
|---|---|---|---|---|
| ***Sporobolus cryptandrus*** | | **13.40±1.33** | **34.00±1.78** | **67.30±3.58** |
| *Alkanna tinctoria* | | | 0.01±0.01 | |
| *Alyssum tortuosum* | | | | 0.06±0.05 |
| *Ambrosia artemisiifolia* | 1.20±0.63 | 0.43±0.20 | 2.75±0.55 | 0.48±0.21 |
| *Anthemis ruthenica* | | | 0.05±0.05 | |
| *Arenaria serpyllifolia* | | 0.02±0.01 | | 0.07±0.05 |
| *Artemisia campestris* | | | | 0.05±0.05 |
| *Asclepias syriaca* | 0.55±0.50 | | | |
| *Ballota nigra* | 0.50±0.50 | | | |
| *Bassia laniflora* | 0.13±0.07 | 0.25±0.13 | 0.57±0.20 | 0.82±0.33 |
| *Bothriochloa ischaemum* | 1.00±1.00 | 4.71±2.12 | 0.05±0.05 | 0.80±0.80 |
| *Bromus squarrosus* | 0.25±0.20 | 0.34±0.14 | 0.07±0.05 | 0.08±0.05 |
| *Bromus tectorum* | | 0.40±0.31 | 0.35±0.30 | 0.72±0.33 |
| *Calamagrostis epigejos* | 6.60±4.92 | | | |
| *Carex liparicarpos* | 7.75±2.80 | 7.65±3.54 | 1.70±0.30 | 1.70±0.55 |
| *Carex stenophylla* | | 0.05±0.05 | | |
| *Cenchrus spinifex* | | | 0.01±0.01 | |
| *Centaurea arenaria* | 0.10±0.10 | 1.30±0.40 | 3.21±1.74 | 0.67±0.42 |
| *Chondrilla juncea* | | | | 0.10±0.10 |
| *Convolvulus arvensis* | 1.00±1.00 | | | |
| *Conyza canadensis* | 0.20±0.15 | 0.05±0.05 | 0.55±0.26 | 0.56±0.26 |
| *Crataegus monogyna* | | | 0.70±0.70 | 0.40±0.40 |
| *Crepis foetida* subsp. *rhoeadifolia* | 2.01±2.00 | 0.41±0.16 | 1.56±0.41 | 1.80±0.95 |
| *Cynodon dactylon* | 2.06±1.52 | 0.40±0.30 | | 0.30±0.30 |
| *Echium vulgare* | 2.00±2.00 | | | |
| *Elymus repens* | 12.46±8.24 | | | |
| *Eryngium campestre* | 0.40±0.23 | 0.61±0.26 | 0.10±0.07 | 0.50±0.26 |
| *Erysimum diffusum* | 0.01±0.01 | 0.01±0.01 | | |
| *Euphorbia cyparissias* | 3.93±1.79 | 2.70±1.96 | 0.10±0.10 | 0.40±0.40 |
| *Euphorbia seguieriana* | 0.44±0.30 | 5.65±1.97 | 1.80±0.55 | 1.51±0.76 |
| *Festuca vaginata* | 3.00±3.00 | 9.40±5.17 | 5.70±3.46 | 0.60±0.34 |
| *Festuca wagneri* | | 1.30±0.60 | | |
| *Galium verum* | 1.75±0.98 | | | |
| *Koeleria glauca* | 0.30±0.30 | 0.10±0.10 | | 0.30±0.21 |
| *Koeleria pyramidata* | | 0.35±0.30 | 0.10±0.10 | |
| *Linaria genistifolia* | | 0.10±0.10 | 0.20±0.20 | |
| *Medicago falcata* | 0.22±0.20 | 0.10±0.10 | | |
| *Medicago minima* | 0.01±0.01 | 0.16±0.10 | 0.26±0.11 | 0.17±0.10 |
| *Minuartia glomerata* | | 0.21±0.13 | 0.40±0.30 | 0.05±0.05 |
| *Poa angustifolia* | 5.70±3.95 | | | 0.10±0.10 |
| *Poa bulbosa* | | 0.11±0.10 | 0.01±0.01 | 0.14±0.06 |
| *Polygonum arenarium* | 0.03±0.02 | 0.58±0.19 | 0.57±0.21 | 0.14±0.10 |

**Electronic Appendix 2E continued.**

| Species | I | II | III | IV |
|---|---|---|---|---|
| *Potentilla incana* | 6.00±3.01 | 3.30±0.94 | | 1.00±1.00 |
| *Robinia pseudoacacia* | | | 0.40±0.40 | |
| *Salsola kali* | | 0.10±0.10 | 0.05±0.05 | 0.02±0.01 |
| *Scirpoides holoschoenus* | | 0.31±0.30 | 0.10±0.10 | |
| *Secale sylvestre* | | 0.30±0.21 | 0.01±0.01 | 0.20±0.11 |
| *Setaria viridis* | 0.08±0.06 | 0.73±0.51 | 0.07±0.05 | 0.10±0.10 |
| *Silene conica* | | 0.01±0.01 | 0.02±0.01 | 0.11±0.07 |
| *Silene latifolia* subsp. *alba* | 2.10±1.99 | | | |
| *Stipa capillata* | | 0.30±0.21 | | |
| *Stipa pennata* | 12.80±4.39 | 3.50±2.42 | 1.20±0.49 | 0.70±0.70 |
| *Teucrium chamaedrys* | 8.50±5.97 | | | |
| *Thymus pannonicus* | 4.05±2.50 | 4.51±2.76 | 1.21±0.68 | 0.51±0.34 |
| *Tragopogon orientalis* | | 0.05±0.05 | | |
| *Verbascum lychnitis* | 0.10±0.10 | 0.10±0.10 | 0.20±0.20 | |
| *Veronica prostrata* | 0.10±0.10 | | | |
| *Viola rupestris* | 0.05±0.05 | | | |
