## Supplementary material for "Sand dropseed (*Sporobolus cryptandrus*) – A new pest in Eurasian sand areas?": E-Appendix3

**Electronic Appendix 3.** Seed bank composition of the sites with an increasing cover of *Sporobolus cryptandrus*. In the table seedling numbers are shown.

| | | **Sporobolus cover in the vegetation** | | | | | | | | | | | | | | | |
|---|---|---|---|---|---|---|---|---|---|---|---|---|---|---|---|---|---|
| | | **I: 0%** | | | | **II: 1-25%** | | | | **III: 26-50%** | | | | **IV: 50-75%** | | | |
| **Species in the seed bank** | | **0-2.5** | **2.5-5** | **5-7.5** | **7.5-10** | **0-2.5** | **2.5-5** | **5-7.5** | **7.5-10** | **0-2.5** | **2.5-5** | **5-7.5** | **7.5-10** | **0-2.5** | **2.5-5** | **5-7.5** | **7.5-10** |
| *Arenaria leptoclados* | **502** | 108 | 24 | 11 | 10 | 238 | 31 | 24 | 23 | 8 | 7 | 2 | 2 | 3 | 7 | 3 | 1 |
| *Portulaca oleracea* | **492** | 12 | 57 | 34 | 39 | 54 | 45 | 19 | 22 | 23 | 35 | 35 | 10 | 24 | 48 | 19 | 16 |
| ***Sporobolus cryptandrus*** | **320** | 3 | 22 | 8 | 9 | 39 | 4 | 3 | 6 | 70 | 26 | 9 | 5 | 90 | 13 | 8 | 5 |
| *Potentilla argentea* | **200** | 28 | 2 | 14 | 8 | 31 | 16 | 9 | 2 | 20 | 10 | 10 | 4 | 21 | 15 | 8 | 2 |
| *Digitaria sanguinalis* | **153** | 29 | 2 | 2 | 8 | 88 | 5 | 7 | 5 | 4 | | 2 | | 1 | | | |
| *Cerastium semidecandrum* | **104** | 16 | 3 | 3 | 2 | 60 | 6 | 1 | 2 | 3 | 1 | | | 7 | | | |
| *Veronica* sp. | **65** | 19 | 11 | 2 | 8 | 4 | 4 | 2 | 1 | 6 | 3 | 1 | 1 | 2 | 1 | | |
| *Poa annua* | **55** | 6 | 1 | | | 9 | | | | 24 | 1 | 1 | | 12 | | | 1 |
| *Scleranthus annuus* | **43** | 9 | 1 | | | 25 | | | | | | | | 8 | | | |
| Poaceae sp. | **37** | 3 | | 2 | | 3 | | | | 15 | 3 | 1 | | 9 | | 1 | |
| *Capsella bursa-pastoris* | **34** | 20 | 1 | 3 | 7 | 1 | | | | | | | | 2 | | | |
| *Erophila verna* | **29** | 3 | 1 | | | 5 | 2 | | 1 | 5 | | | | 12 | | | |
| *Erodium cicutarium* | **23** | | | | | 2 | | | | 16 | | 1 | 1 | 3 | | | |
| *Plantago lanceolata* | **15** | 10 | 3 | | | 2 | | | | | | | | | | | |
| *Oxalis corniculata* | **10** | 0 | 2 | 3 | | 1 | 1 | | | | 1 | | 2 | | | | |
| *Conyza canadensis* | **8** | 7 | 1 | | | | | | | | | | | | | | |
| *Bromus* cf. *arvensis* | **5** | | | | | 4 | | | | 1 | | | | | | | |
| *Eragrostis minor* | **5** | | | | | 1 | | | | | | | | 1 | 2 | | 1 |
| *Medicago minima* | **5** | 1 | | 1 | 1 | | 1 | | | | | | | 1 | | | |
| *Carex* cf. *stenophylla* | **4** | | | | | 1 | | 1 | | 1 | | | | | | 1 | |
| *Poa angustifolia* | **4** | | | | | | | | | | | | | 2 | 2 | | |
| *Epilobium* sp. | **3** | 1 | | | | 2 | | | | | | | | | | | |
| *Juncus articulatus* | **2** | 1 | | | | | | | 1 | | | | | | | | |
| *Medicago lupulina* | **2** | | | 1 | | | | | | | | 1 | | | | | |
| *Melandrium album* | **2** | | | | | | | | | | | 2 | | | | | |
| *Trifolium repens* | **2** | 2 | | | | | | | | | | | | | | | |
| *Stellaria media* | **2** | | | | | 2 | | | | | | | | | | | |
| *Vicia lathyroides* | **2** | | | | | | | | | 2 | | | | | | | |
| *Eragrostis minor* | **1** | | | | | | | 1 | | | | | | | | | |

**Electronic Appendix 3.** Continued.

| | | Sporobolus cover | | | | | | | | | | | | | | | |
|---|---|---|---|---|---|---|---|---|---|---|---|---|---|---|---|---|---|
| | | **0%** | | | | **1-25%** | | | | **26-50%** | | | | **50-75%** | | | |
| **Species** | | **0-2.5** | **2.5-5** | **5-7.5** | **7.5-10** | **0-2.5** | **2.5-5** | **5-7.5** | **7.5-10** | **0-2.5** | **2.5-5** | **5-7.5** | **7.5-10** | **0-2.5** | **2.5-5** | **5-7.5** | **7.5-10** |
| *Poa pratensis* | **1** | | | | | 1 | | | | | | | | | | | |
| *Juncus effusus/conglomeratus* | **1** | 1 | | | | | | | | | | | | | | | |
| *Anthemis ruthenica* | **1** | 1 | | | | | | | | | | | | | | | |
| ***Total number of seedlings*** | **2132** | **280** | **131** | **84** | **92** | **573** | **115** | **67** | **63** | **198** | **87** | **65** | **25** | **198** | **88** | **40** | **26** |

**Footnote:** The seed density for 1 $m^2$ can be calculated if the above scores are multiplied with ~26.53 (surface area of 30 soil cores in total from 3 plots, 4 cm diameter of a cylindrical corer compared to 10,000 $cm^2$). This means for example that the seed density of *Arenaria leptoclados* for I 0-2.5cm layer (108 seeds) reads as 2,865 seeds/$m^2$.
